# Modeling predicts a biochemical feedback mechanism underlying astrocyte calcium refractory periods

**DOI:** 10.64898/2026.01.12.699127

**Authors:** James Cheng Peng, Katharina Merten, Mary Kaye Duff, Nicholas Nelson, Yizhi Wang, Daniela Duarte, Yue Wang, Lin Tian, Axel Nimmerjahn, Guoqiang Yu

## Abstract

Astrocytes exhibit intracellular calcium fluctuations in response to neuronal activity. Repeated receptor stimulation can induce a transient suppression of calcium signaling—a refractory period—yet the underlying mechanisms and timescales of this phenomenon during behavior remain poorly understood. Here, we present a biophysically grounded computational model of astrocytic calcium signaling that incorporates a novel feedback mechanism mediated by conventional protein kinase C (cPKC) and predicts the refractory phenomenon. Unlike previous models developed in vitro, our model is directly validated using *in vivo* two-photon calcium imaging data from behaving mice. It closely recapitulates astrocytic calcium dynamics across both time and frequency domains, including the emergence of refractory periods and their negative correlation with inter-stimulus intervals. Simulations further predict the timing of recovery from refractory states, consistent with experimental observations. This work provides a mechanistic explanation for astrocytic refractory behavior and establishes a framework for integrating computational modeling with *in vivo* functional imaging.

## INTRODUCTION

Astrocytes, a type of non-neuronal cell in the central nervous system, play essential roles in maintaining molecular, cellular, and network homeostasis, and are directly involved in brain information processing^1,2^. Unlike neurons, which generate large shifts in membrane potential upon activation, astrocytes exhibit limited electrical excitability^3^. Instead, they show excitability to diverse extracellular stimuli such as neurotransmitters, neuromodulators, and other signaling molecules^4,5^ through intracellular calcium transients with various patterns^1^. These calcium signals trigger diverse downstream pathways and can have profound impacts on synaptic transmission, circuit activity, and behavior^6^. However, how astrocytes integrate the multifactorial inputs and mediate the downstream effects is poorly understood. A particularly intriguing feature of astrocytic calcium signaling is the so-called *refractory period* —a transient suppression of calcium responses following large transients, during which reactivation is inhibited. This phenomenon has been observed and replicated across multiple studies and brain regions^7–12^, yet its mechanistic basis remains unclear. Understanding the origin of this refractory effect is critical because it defines the intrinsic timescale over which astrocytes can engage with neural circuits, constraining how they integrate successive inputs and modulate synaptic and network activities.

Computational modeling of astrocyte calcium dynamics has emerged as a powerful approach for dissecting the mechanisms that generate refractory periods, as it formalizes the complex interplay between extracellular cues and intracellular signaling pathways and enables the systematic testing of mechanistic hypotheses that are difficult to isolate experimentally. Early models, such as the De Young–Keizer and Li–Rinzel formulations^13,14^, successfully captured inositol 1,4,5trisphosphate (IP_3_)-dependent calcium-induced calcium release (CICR) but did not incorporate upstream signaling cascades or astrocyte-specific regulatory mechanisms. Subsequent modeling efforts introduced receptor-driven IP_3_ production, as exemplified by the G-ChI framework^15^, and incorporated voltage-gated calcium channels (VGCCs) to better reflect the diversity of calcium influx mechanisms^16^.

Later generations of models integrated morphological complexity and stochastic signaling features to represent spatially heterogeneous dynamics more accurately^17–20^, while more recent approaches have extended these frameworks to describe long-range calcium wave propagation^21,22^. Network- and multi-compartment-level models have further expanded this work by incorporating bidirectional neuron–astrocyte interactions and emergent network dynamics^22–27^ Additionally, newer models have begun to account for neuromodulatory and pathological influences, including dopamine- and A*β*-dependent modulation of astrocyte calcium signaling^28–30^, thereby bridging intracellular dynamics with broader physiological and disease contexts.

However, most existing astrocyte calcium models have been developed and validated primarily against cell-culture or acute brain-slice data and are typically tailored to a limited set of input pathways matched to those preparations, rather than to the diverse, behaviorally derived signals that drive astrocyte activity *in vivo*. Yet, astrocyte calcium dynamics *in vivo* differ fundamentally from those measured under such conditions^2,31,32^, limiting the utility of these models for decoding the complex relationship between astrocytic activity and behavior in awake animals. In particular, none of the current modeling approaches can explain or predict the refractory period described above.

At first glance, the focus on simplified, culture-based models may seem justified, as modeling astrocyte calcium dynamics *in vivo* would require accounting for a far greater diversity of signaling molecules, modulatory inputs, and behavioral states, an endeavor that appears prohibitively complex. However, when viewed from a data-driven perspective, this apparent complexity can be transformed from a liability into an asset. By designing experimental paradigms that systematically elicit and record diverse behavioral conditions, the resulting data richness can be leveraged to train models using machine learning approaches, thereby enabling the prediction of astrocytic calcium dynamics under physiologically relevant conditions.

Here, we developed a comprehensive model of astrocytic calcium signaling, effectively a “virtual astrocyte,” that (1) integrates not only neuromodulatory but also synaptic inputs with cytosolic, endoplasmic reticulum (ER), and mitochondria calcium dynamics, (2) involves more than 30 molecules and ions and accounts for the dynamic interactions between them, (3) is directly tested and validated using *in vivo* calcium imaging data from awake, behaving mice. An enabling factor of this study is the design of a behavior paradigm, which allows multiple aspects of the behavior and an array of outcomes to be recorded simultaneously. The behavioral parameters, such as running speed, reward, and visual stimulus delivery, were used to approximate the neuromodulatory and synaptic inputs. We used machine learning techniques to optimize the 98 unknown parameters based on over 1000 quantitatively different trials, where the diversity of responses effectively prevents overfitting even for a model considering 30 molecules and ions.

Our computational analysis and modeling enabled us to test and ultimately refute the long-standing but previously untested ER depletion hypothesis^7,8,10,11^, which posited that depletion of endoplasmic reticulum calcium stores underlies the astrocytic refractory period. Both our empirical observations and mathematical modeling demonstrated that depletion cannot account for this phenomenon. The high resting calcium concentration within the ER (hundreds of micromolars) relative to the cytosol (hundreds of nanomolars), combined with the ER’s substantial volume (15–70% of the cytoplasm), makes complete depletion highly improbable^33–36^. This conclusion was further supported by statistical analyses showing that, when conditioned on the interval between two consecutive calcium events, the amplitude of the preceding event did not influence the magnitude of the subsequent one.

Our modeling further identified a negative feedback loop mediated by conventional protein kinase C (cPKC), which is activated during calcium transients by diacylglycerol (DAG) and calcium and, in turn, phosphorylates and inhibits upstream Gq-GPCR and phospholipase C (PLC) signaling^15,35,37–39^. The model also predicted that cPKC activity remains elevated for a defined temporal window following activation, thereby suppressing subsequent calcium events and providing a mechanistic explanation for the refractory period.

Beyond explaining the refractory period, our modeling framework accurately reproduced astrocytic calcium dynamics across both time and frequency domains. The amplitude of calcium responses scaled positively with the strength of neuromodulatory and synaptic inputs, and the model revealed the importance of nonlinear integration, whereby simultaneous neuromodulatory and synaptic signaling produced synergistic amplification of calcium signals. When behavioral trials were categorized as rewarded or non-rewarded, the model successfully captured the overall trend of enhanced responses during rewarded trials. However, it tended to overestimate peak amplitudes in rewarded conditions and underestimate them in non-rewarded ones, suggesting the existence of additional, unmodeled signaling pathways or intracellular buffering mechanisms Finally, beyond astrocytic calcium signaling per se, our work establishes a generalizable framework for constructing virtual cells—models that integrate behaviorally informed experimental paradigms with data-driven parameter inference to achieve biophysically grounded mechanistic models of cellular dynamics.

## RESULTS

### Biophysical and molecular architecture of the astrocyte calcium model

We developed an end-to-end computational framework that links animal behavior, behavior-derived extracellular molecular inputs, and astrocytic intracellular calcium transients, while capturing intrinsic features of astrocytic dynamics such as refractory periods.

In our experiments, mice performed a visual detection task while astrocytic calcium activity in the motor cortex was monitored using two-photon microscopy. The two parallel pipelines underlying this framework are illustrated schematically in Fig. 1A. In the data-driven approach (Fig. 1A, top), *in vivo* calcium imaging data are preprocessed to extract astrocytic calcium traces and event timings. In the model-driven approach (Fig. 1A, bottom), behavioral events such as locomotion, sensory cues, and reward delivery are transformed into behavior-derived molecular input signals, which drive the computational model to generate predicted calcium dynamics. Together, these pipelines bridge experimental measurements with physiologically grounded simulations of astrocyte signaling.

**Figure 1.**
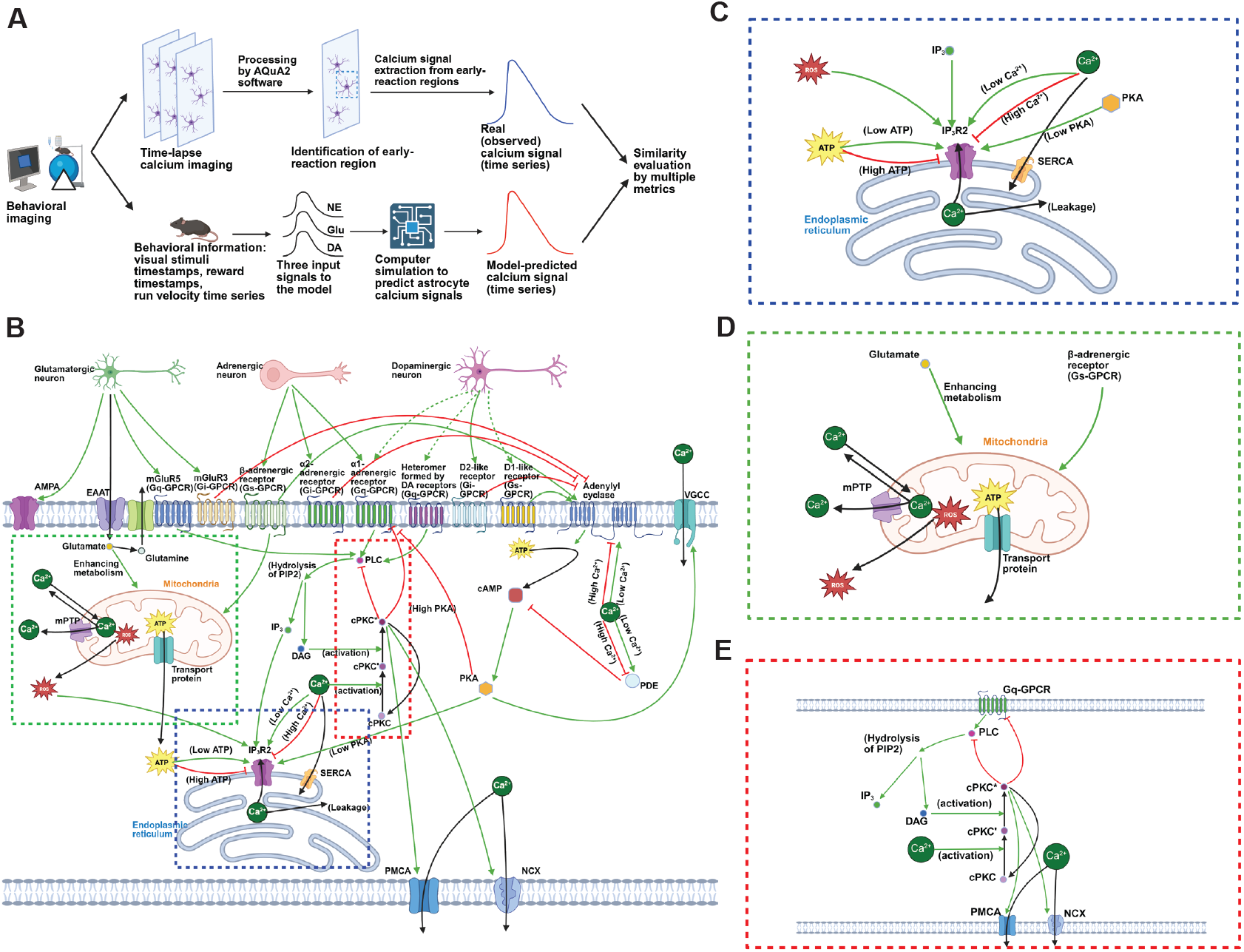
Schematic of analysis pipelines and modeled astrocytic calcium signaling pathways. **(A)** Computational workflow for calcium signal extraction and prediction. Experimental calcium events were extracted from imaging data (top), while model-predicted signals were generated from trial events and associated molecular inputs (bottom). **(B)** Overview of the signaling pathways incorporated into the model, corresponding to the computer simulation part in (A). Molecular inputs (top) include glutamate, NE, and DA, which act on receptors and transporters at the plasma membrane. The model output is the resultant intracellular calcium dynamics. Green and red arrows indicate activation and inhibition, respectively; black arrows denote molecular transport or conversion; dashed arrows mark pathways that were included but rendered functionally inactive or excluded. For example, due to the low expression of D1-like receptors in cortical astrocytes^40^, pathways involving D1-like receptors (alone or as D1/D2 heteromers) were assigned a coefficient of zero. Similarly, while DA has been reported to act on *α*1-adrenergic receptors in the prefrontal cortex^41^, this pathway was excluded here. **(C–E)** Magnified views of the blue, green, and red regions in (B), illustrating: (C) IP_3_-mediated CICR from the ER, (D) mitochondria-buffered calcium signaling, and (E) a cPKC-mediated feedback mechanism that inhibits Gq-GPCR and PLC activity, contributing to regulation of the astrocytic calcium refractory period. Figures created with BioRender.

Fig.1B summarizes the intracellular pathways incorporated into the model-driven pipeline (see also Table1 and Appendix S6). Astrocytic calcium signaling is shaped by diverse neurotransmitters and neuromodulators and involves nonlinear interactions between these pathways^2,34,42^. We focused on three key inputs: (1) glutamate, a fast-acting transmitter; (2) norepinephrine (NE); and (3) dopamine (DA), both slower neuromodulators. For intracellular signaling, we emphasized pathways directly involved in calcium regulation for computational tractability and to enable comparisons between simulated and experimental calcium dynamics.

Behavioral events, including locomotion (run velocity), sensory cues, and reward delivery, were used to infer glutamate, NE, and DA input signals (Fig. 2A-C). Activation of the corresponding astrocytic receptors initiates intracellular cascades that collectively generate calcium transients Our model incorporates: (i) IP_3_-mediated calcium-induced calcium release (CICR) through IP_3_R2 (Fig. 1C); (ii) the calcium–cAMP–PKA circuit; (iii) mitochondrial calcium buffering (Fig. 1D); and (iv) calcium flux across the plasma membrane via voltage-gated calcium channels (VGCCs), AMPA receptors, plasma membrane calcium-ATPase (PMCA), and sodium-calcium exchangers (NCX) We also included a negative feedback loop mediated by conventional protein kinase C (cPKC) (Fig. 1E). Among IP_3_ receptor isoforms, IP_3_R2 was chosen given its predominant expression in mouse cortical astrocytes^40^. AMPA receptors, despite being largely calcium-impermeable in most brain astrocytes, were included for generalizability, as calcium-permeable subtypes exist in specific subpopulations such as olfactory bulb astrocytes and cerebellar Bergmann glial cells^43,44^ Together, these pathways transform behavior-driven molecular inputs into intracellular calcium signals that capture both amplitude and refractory properties of astrocytic activity.

**Figure 2.**
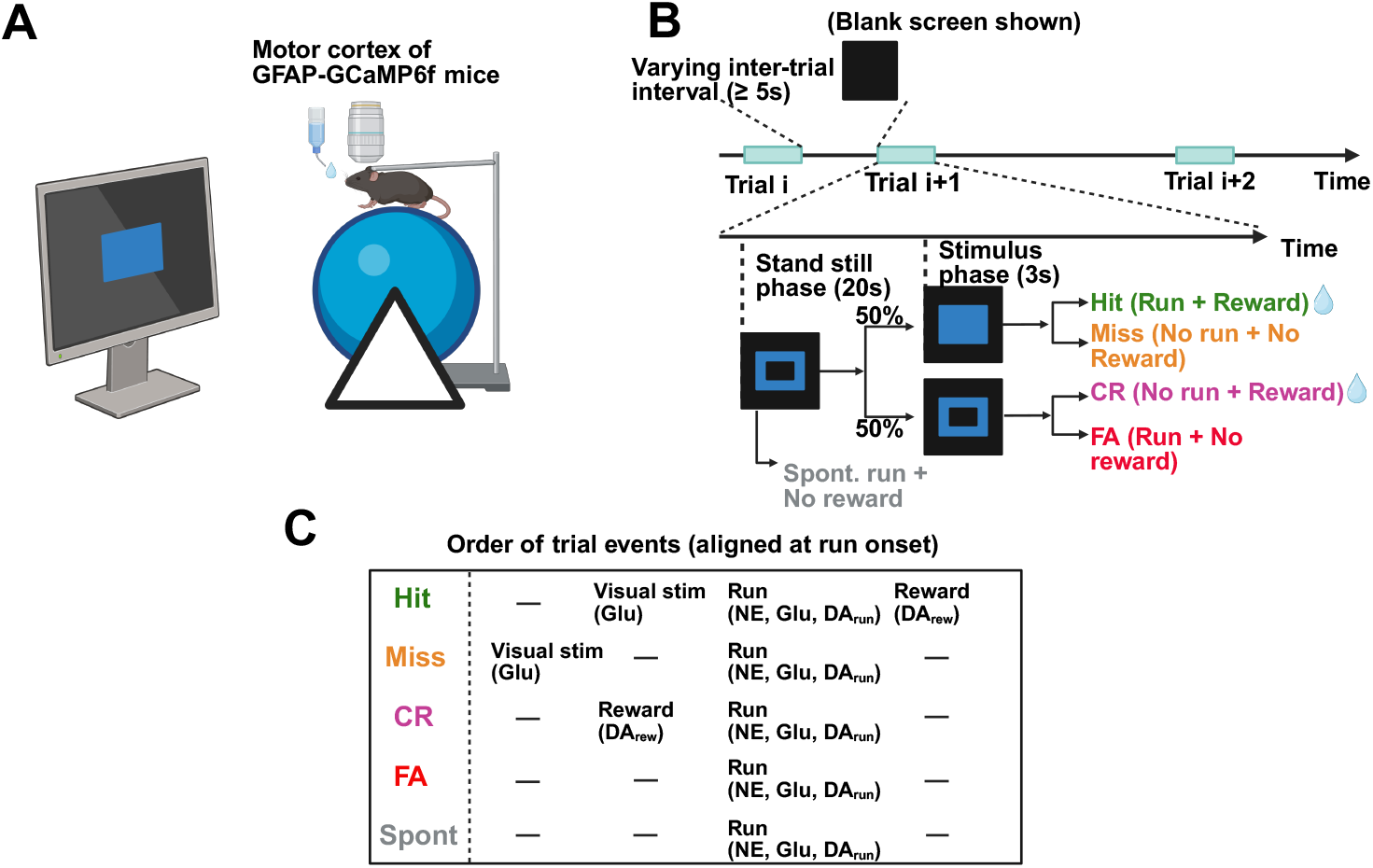
Experimental workflow for behavioral imaging. **(A)** Schematic of the experimental setup. Head-fixed GFAP-GCaMP6f mice were placed on a spherical treadmill facing a monitor that presented visual stimuli to guide behavior. Astrocytic calcium activity in the motor cortex was recorded using two-photon imaging while the trained animals performed the visual detection task. **(B)** Schematic of the behavioral protocol (bottom) and trial sequence (top). Trials began when mice stopped running for 1 s. An open blue square signaled mice to remain stationary for 20 s. After this period, either a filled blue square (50% of trials) appeared or the open blue square remained displayed for 3 s. If the filled square appeared, mice had to start running within the 3-s window to obtain a water reward (“Hit”); failure to run was scored as a “Miss.” If no filled square appeared, mice had to remain stationary for 3 s to receive a reward (“CR”); in contrast, running in this case was scored as a “FA.” Spontaneous running during the initial 20-s stationary period aborted the trial (“Spont”). After an inter-trial interval of ≥5 s, mice could initiate the next trial. The filled square was presented at two intensities—salient or threshold (see **STAR METHODS** for details). **(C)** Table summarizing the sequence of events and putative molecular signals for each trial type, aligned to run onset. Figures created with BioRender.

The IP_3_-mediated CICR pathway is a well-characterized component of astrocytic calcium signaling (Fig. 1C). Gq-GPCR activation stimulates PLC, which hydrolyzes PIP_2_ into IP_3_. IP_3_ binds to ER-localized IP_3_Rs, triggering calcium release into the cytoplasm. IP_3_R activity is shaped by multiple intracellular factors: IP_3_ (positive regulation), calcium and ATP (bell-shaped modulation), as well as reactive oxygen species (ROS) and PKA (enhancers)^45–48^.

Astrocytes integrate Gq-GPCR inputs synergistically. For example, glutamate not only generates IP_3_ but also fuels metabolic pathways, increasing ATP production, which further enhances IP_3_R activity. Co-activation by glutamate and NE can therefore yield amplified calcium responses compared to either input alone^49^. ROS, modeled as a linear function of glutamate-driven mitochondrial activity, and PKA, modeled through the cAMP-PKA cascade, provide additional modulation.

We placed particular emphasis on ATP, given the bell-shaped relationship between ATP levels and IP_3_R opening rates^45^. Because ATP was not directly measured *in vivo*, we modeled it as a proxy signal derived from glutamate and NE inputs—reflecting glutamate’s role in mitochondrial ATP synthesis and NE’s link to locomotor energy demand. This modeled ATP reflects inferred production rather than absolute concentrations. Reported in situ astrocytic ATP levels (0.7–1.3 mM)^50^ support its facilitative role in IP_3_R2 activity.

We developed a mathematical formulation of IP_3_R activity based on channel opening rate data^45^, optimized parameters via gradient descent, and incorporated modulatory effects of ROS and PKA through Hill functions. Complete formulations are given in Appendix S2.

Under resting conditions, mitochondrial calcium is assumed to approximate cytosolic levels but rises 10–20 fold upon stimulation. Influx was modeled as a Hill function of cytosolic calcium, while efflux depended on both ROS levels and the mitochondrial–cytoplasmic calcium gradient (Fig. 1D; see also **STAR METHODS**)^51^.

We also explicitly modeled the calcium influx to cytoplasm via VGCCs and AMPA receptors, and included the effects of calcium extrusion via PMCA and NCX in the model of calcium efflux (Appendix S1, Eqs. (11), (24), (28), (30)). Additionally, activated by calcium and DAG, cPKC translocates to the plasma membrane, where it inhibits Gq-GPCR and PLC activity, thereby dampening calcium signaling^15,35,37,38^. PKC-mediated feedback (Fig. 1E) has been implicated in terminating calcium spikes and modulating interspike intervals in astrocytes, though its physiological role *in vivo* remains unresolved. Here, we include it as a candidate mechanism underlying astrocytic refractory periods following repeated stimuli such as locomotion.

The overall system was implemented as a set of ordinary differential equations, assuming a mean-field approximation in which each compartment is considered well mixed. Each equation describes the temporal change of a chemical species, receptor state, or calcium flux between the extracellular space, cytoplasm, ER, and mitochondria. Details of the equations are given in Appendix S1; parameter optimization is described in Appendix S4.

### Behavior-derived molecular inputs included in the model

Due to current technical limitations, calcium, glutamate, NE, and DA cannot be measured simultaneously using two-photon microscopy. We therefore inferred transmitter release from behavioral readouts, based on well-established links between behavior and activation of corresponding molecular pathways^49,52–55^.

NE release is closely linked to locomotion and drives widespread astrocytic calcium activity In our model, NE is represented as a leaky integrator of run velocity, an approach supported by recent work showing that leaky integration of behavioral variables (e.g., paw movement, pupil diameter) effectively captured correlations with astrocytic calcium signals in the hippocampus^12^ NE has also been shown to correlate with cortical astrocytic calcium activity^52^. Thus, leaky integration of run velocity provides a biologically plausible intermediary between locomotion and astrocytic calcium dynamics. In our model, NE acts via *α*1-adrenergic receptors (Gq-coupled GPCRs) that drive calcium signaling through the IP_3_ pathway, *β*-adrenergic receptors (Gs-coupled GPCRs), and *α*2-adrenergic receptors (Gi-coupled GPCRs), which regulate adenylyl cyclase, cAMP signaling, and metabolism (Fig. 3A, top).

**Figure 3.**
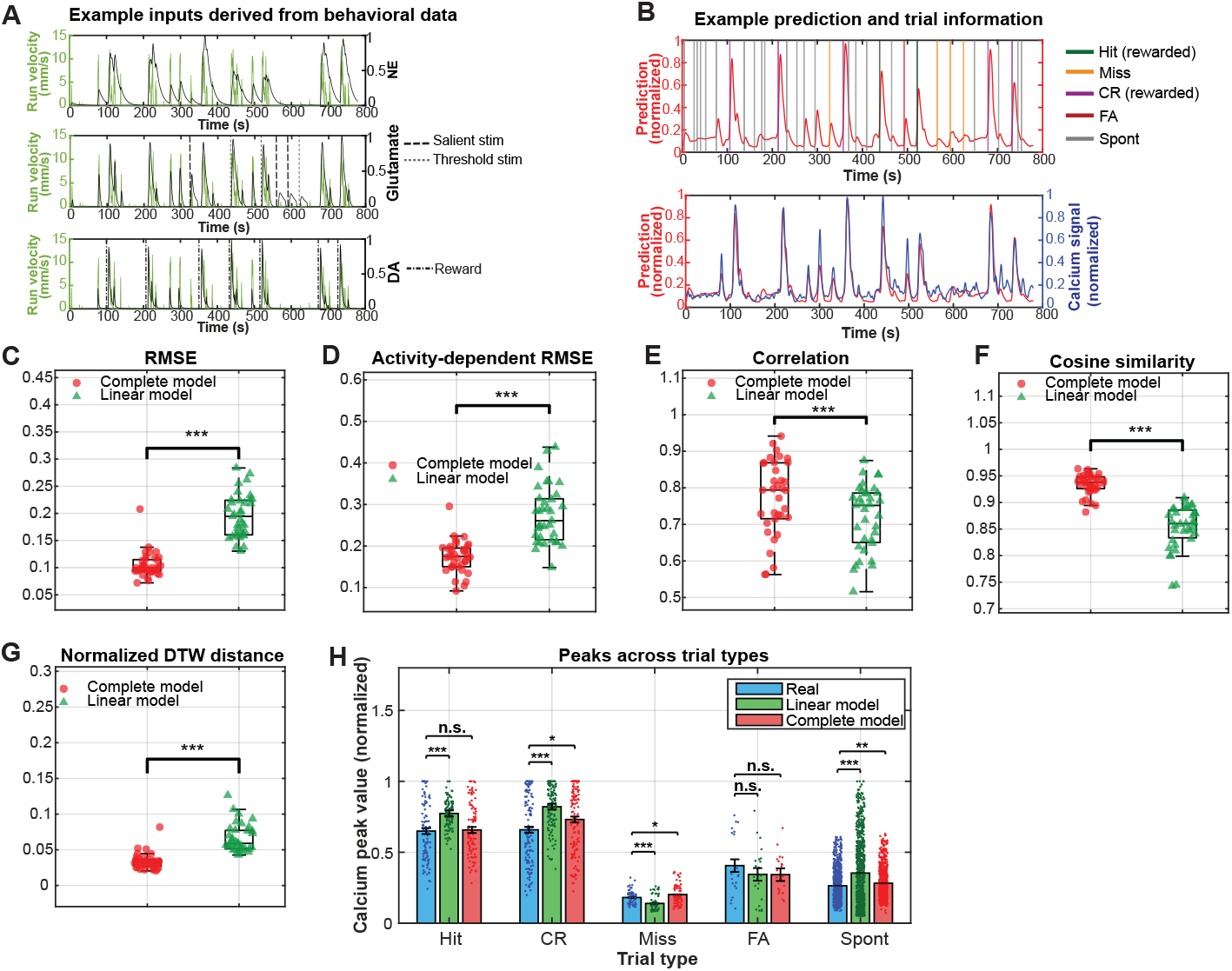
The model predicts trial-specific astrocytic calcium responses from behavior-derived molecular inputs. **(A)** Example recording showing three molecular input signals derived from run velocity, visual stimuli, and reward. NE (top) was modeled as a leaky integrator of run velocity (green curves). Glutamate (center) was modeled as impulses triggered by running and visual stimuli. DA (bottom) was modeled as impulses triggered by running and rewards. Vertical lines indicate stimulus and reward onset (salient: dashed; threshold: dotted; reward: dash-dotted). **(B)** Comparison of model-predicted (red) and real calcium signals (blue) for the example recording in (A). The two signals (bottom) show close correspondence. Vertical colored lines mark different trial types aligned at run onset. **(C-G)** Box plots showing, for each of the five performance metrics, the comparison between the complete model and the linear model across all 35 recording sessions. Each metric quantifies how well each model’s predicted calcium signal matches the real calcium signal obtained from imaging data. Lower values indicate better performance for RMSE, activity-dependent RMSE, and normalized DTW distance, whereas higher values indicate better performance for correlation and cosine similarity. Paired t-tests comparing the two models yielded the following p-values: RMSE (3.28 × 10^−19^), activity-dependent RMSE (6.74 × 10^−10^), correlation (2.98 × 10^−5^), cosine similarity (2.74 × 10^−14^), and normalized DTW distance (7.02 × 10^−15^). **(H)** Trial-based peak amplitudes comparison among the real calcium signals (blue), signals predicted by the linear model (green), and signals predicted by the complete model (red). The bar plot shows mean calcium peak values (±SEM) across the five trial types: Hit (*n* = 100, real: 0.649 ± 0.021, linear: 0.773 ± 0.010, complete: 0.657 ± 0.018), CR (*n* = 135, real: 0.660 ± 0.021, linear: 0.821 ± 0.013, complete: 0.731 ± 0.020), Miss (*n* = 71, real: 0.182 ± 0.006, linear: 0.140 ± 0.006, complete: 0.201 ± 0.008), FA (*n* = 22, real: 0.410 ± 0.045, linear: 0.344 ± 0.041, complete: 0.342 ± 0.029), and Spontaneous (*n* = 728, real: 0.270 ± 0.005, linear: 0.353 ± 0.009, complete: 0.281 ± 0.004). n.s. p-value ≥ 0.05; * p-value*<* 0.05; ** p-value*<* 0.01; *** p-value*<* 0.001. Error bars represent SEM. Colored dots in (C-G) denote individual recordings, and in (H) denote individual trials. measures.

DA release in the cortex is associated with both reward and locomotion. For example, electrical stimulation of dopaminergic inputs from the ventral tegmental area to the motor cortex reinforces locomotor behavior and enhances reward-driven performance^54^. Accordingly, DA inputs in our model are triggered by reward or run onset (Fig. 3A, bottom). Each DA impulse is modeled as the product of two Hill functions (Appendix S1, Eq. (13)), capturing the rapid rise and slower decay observed in two-photon imaging of the fluorescent dopamine indicator dLight1.2 in motor cortex^55,56^. DA signaling is implemented through heteromeric receptor complexes (Gq-coupled), D1-like receptors (Gs-coupled GPCRs), and D2-like receptors (Gi-coupled GPCRs), which regulate calcium signaling indirectly via the cAMP-PKA pathway^48^. Although D1-like receptor expression is low in mouse cortical astrocytes^40,57^, they were included to allow model extension to other astrocyte populations. Users can tune receptor-specific coefficients during model fitting to reflect region- or cell–type–specific gene expression patterns.

Extracellular glutamate levels increase in the cortex during locomotion and are further modulated by sensory inputs^53,58^. We therefore modeled glutamate release as impulses triggered by running and sensory stimuli (Fig. 3A, center). Each impulse is represented by the product of two Hill functions (Appendix S1, Eq. (15)), producing a sharp rise and slower decay. Although the mathematical form differs from NE integration, the resulting signal traces show similar transient dynamics (Fig. 3A). In astrocytes, glutamate acts through metabotropic and ionotropic receptors as well as transporters^59^. Among metabotropic receptors, mGluR5 (Gq-coupled) can evoke calcium release via IP_3_-mediated CICR^15^. While not typically expressed in adult cortical astrocytes under physiological conditions^60^, mGluR5 was included to allow applications involving developmental or pathological (re-)expression^61^. mGluR3 (Gi-coupled) is expressed in mouse astrocytes and regulates cAMP and metabolism. Among ionotropic receptors, NMDA receptors were excluded due to low astrocytic expression^40,57^, whereas AMPA receptors were included given their high expression^40,57^. Since most AMPA receptors are calcium-impermeable^62^, but some astrocytic subtypes (e.g., in olfactory bulb and cerebellar cortex) are calcium-permeable^43,44^, we assigned a small coefficient to represent calcium influx through AMPA receptors, ensuring model generalizability (Appendix S1, Eq. (10)). Finally, glutamate uptake via excitatory amino acid transporters (EAATs) was modeled, reflecting astrocytic roles in glutamate clearance and metabolism^63,64^.

For the Gq-, Gs-, and Gi-coupled GPCR pathways, the effects of the three input molecular signals—NE, DA, and glutamate—were summed to generate the net receptor activation (Appendix S1, Eqs. (21)–(23)). To simulate their combined influence on intracellular signaling, we precomputed the time courses of each input signal. The full model was implemented in MATLAB and numerically integrated using a fourth-order Runge–Kutta method.

### Behavioral and *in vivo* imaging paradigm for model evaluation

To inform and evaluate our model, we performed two-photon calcium imaging in head-fixed mice expressing GCaMP6f in astrocytes (N=4 GFAP-GCaMP6f mice) while they were placed on a spherical treadmill (Fig. 2A). Mice were trained to perform a visual detection task that required either running or remaining still to obtain a reward^9,55^ (Fig. 2B). Recordings in fully trained, expert mice were focused on layer 2/3 of the motor cortex (100-135 *µ*m below the pia), a region engaged during locomotion and previously studied for dopamine signaling^55^.

At the start of each trial, an open blue square appeared after a period of mouse inactivity, signaling the requirement to remain still for 20 seconds. Following this period, either a filled blue square appeared or the open blue square remained displayed. The filled square required the mouse to initiate a run above a threshold velocity and duration to receive a water reward, whereas in its absence, the animal was rewarded if it remained still for an additional 3 seconds. The filled square was shown at either high intensity (salient stimulus) or low intensity (threshold-level stimulus) with equal probability.

Behavioral responses were classified into five trial types: Hit, Miss, Correct Rejection (CR), False Alarm (FA), and spontaneous run (Spont). Hit trials occurred when the mouse ran appropriately in response to the filled square and earned a reward; Miss trials were failures to respond and were not rewarded. CR trials corresponded to correctly remaining still when no filled square was shown and were rewarded, whereas FA trials reflected running in the absence of a filled square and were not rewarded. Spont trials were defined as runs initiated during the 20-second stillness period. Thus, trials could be grouped into two categories: rewarded (Hit, CR) and non-rewarded (Miss, FA, Spont). A schematic overview of the trial structure, aligned to run onset and indicating putative molecular agonists, is provided in Fig. 2C.

Two-photon imaging data were acquired at 30.9 frames per second (fps) and synchronized with behavioral measurements, including run velocity (resampled to the imaging frame rate), visual stimulus (filled square) onset, reward delivery, and inter-trial intervals. Additional methodological details are provided in the **STAR METHODS**.

### Nonlinear mechanisms are essential for accurate calcium signal prediction

Astrocytic calcium signals arise from behavior-driven molecular inputs, and a natural first question is how well these inputs linearly explain the calcium dynamics. We therefore constructed a linear model in which the predicted calcium signal is expressed as a weighted sum of the behavior-derived glutamate, NE, and DA inputs. These three molecular signals are shown for an example recording, illustrating the inputs from which the linear model forms its prediction (Fig. 3A). The weights were optimized to ensure a fair comparison with our full mechanistic model—hereafter referred to as the complete model—which explicitly incorporates nonlinear interactions among inputs as well as the intracellular signaling pathways governing calcium regulation. This makes the linear model a principled benchmark for evaluating the added value of the nonlinear, pathway-resolved structure in the complete model.

To evaluate predictive performance, we compared the linear and complete models across the five metrics used throughout this study—root mean squared error (RMSE), activity-dependent RMSE, correlation, cosine similarity, and normalized dynamic time warping (DTW) distance (Fig. 3C–G). These metrics capture complementary aspects of accuracy and are the same metrics used for parameter optimization of the mechanistic model (Appendix S4). RMSE quantifies the average pointwise deviation across the entire calcium trace, whereas activity-dependent RMSE measures the error specifically during calcium events and therefore reflects performance on physiologically meaningful transients. Correlation assesses temporal synchrony, cosine similarity captures global waveform shape, and normalized DTW distance measures temporal alignment between real and predicted signals. Across all 35 recordings, the complete model consistently outperformed the linear model across every metric.

These findings illustrate the limitations of the linear model and underscore the necessity of nonlinear biochemical mechanisms. Astrocytic calcium signaling is shaped by synergistic interactions among glutamatergic, adrenergic, and dopaminergic pathways, including NE–DA synergy and NE–glutamate synergy^4^. Such nonlinear interactions, along with ATP-dependent modulation, IP_3_R2-mediated calcium-induced calcium release, and history-dependent feedback, cannot be captured by any linear combination of inputs. Our full mechanistic model incorporates these essential nonlinearities, enabling accurate prediction of astrocyte calcium dynamics across diverse behavioral conditions. Together, these results establish the linear model as a principled benchmark and highlight the need to transition to a nonlinear mechanistic framework, which we detail next.

### The model predicts astrocyte calcium dynamics with high accuracy across metrics and trial-type-specific peak amplitudes

Building on the comparison with the benchmark linear model, we next evaluated the predictive performance of the complete mechanistic model in the temporal domain. Fig. 3B shows an example recording, illustrating that the predicted calcium trace closely follows the experimentally measured signal, capturing both the timing and amplitude of major transients. Complementing this qualitative agreement, the model achieved strong performance across the five quantitative metrics—RMSE, activity-dependent RMSE, correlation, cosine similarity, and normalized DTW distance—demonstrating accurate reconstruction of astrocyte calcium dynamics at the whole-trace level (Fig. 3C–G).

While these global similarity metrics are informative, they may obscure trial-type–specific differences in calcium transients. We therefore examined whether the model could reproduce calcium peak amplitudes across individual behavioral trial types, which reflect distinct combinations of glutamatergic and neuromodulatory inputs. Calcium peaks were extracted for Hit, Miss, CR, FA, and Spontaneous trials across all 35 recordings, comprising a total of 1,056 trials. The complete model successfully reproduced the relative ordering of calcium peak amplitudes across trial types, matching the experimentally observed pattern in which Hit and CR trials evoke the largest calcium responses (Fig. 3H). The model also captured the absolute amplitudes of Hit and FA peaks without significant deviation from recorded data.

Some systematic biases were still evident across trial types. The model tended to overestimate peak amplitudes on Miss and CR trials, while producing no significant difference from real data on Hit and FA trials. Spontaneous events showed the largest discrepancy, with predicted peaks consistently lower than the measured values, suggesting contributions from unmodeled behavioral factors—such as fluctuations in attention or arousal, grooming, or whisking—or additional molecular inputs including serotonin^65^ or acetylcholine^66^ (see **DISCUSSION**). Nevertheless, these deviations were markedly smaller than those produced by the linear model: the linear model mis-estimated peak amplitudes in every trial type except FA, with all discrepancies highly significant (p-value *<* 0.001). Overall, the complete model preserved the correct ordering of peak amplitudes across trial types and provided substantially closer agreement with real data than the linear benchmark (Fig. 3H).

Together, these results demonstrate that the nonlinear mechanistic model accurately captures both global temporal features and trial-type–specific modulation of astrocyte calcium dynamics This comprehensive agreement with experimental measurements underscores the necessity of nonlinear biochemical interactions for reproducing *in vivo* astrocytic activity.

### ATP enables synergistic integration of norepinephrine and glutamate inputs

Experimental studies indicate that cortical astrocytes integrate NE and glutamate synergistically during locomotion^49,52^, a behavior associated with increased energy demand and elevated intracellular ATP levels^67^. Although the precise molecular mechanism of NE–glutamate synergy remains unknown, recent evidence indicates that cAMP–PKA signaling is not responsible for this interaction^4^, prompting consideration of alternative pathways. Because ATP modulates IP_3_R2 activity in a bell-shaped manner^45,46^ (Fig. 1C), we hypothesized that locomotion-induced ATP increases facilitate the synergistic integration of NE and glutamate.

To test this, we constructed a reduced model in which ATP levels were held constant rather than variable. By fixing ATP at 0.05 mM (manually optimized for fair comparison with the complete model), we effectively removed ATP-dependent regulation from the system. In this reduced setting, the modulatory influence of ATP on IP_3_R2 is eliminated, and NE–glutamate interactions collapse to a purely additive contribution to calcium signaling. To assess the consequences of ATP removal, we focused on metrics that are sensitive to differences during periods of elevated calcium activity, where ATP-dependent modulation is expected to play the strongest role. Under these metrics, activity-dependent RMSE and normalized DTW distance increased significantly in the ATP-excluded model relative to the complete model (Figure S2A–B). In contrast, whole-trace metrics such as RMSE, correlation coefficient, and cosine similarity, which are dominated by long quiescent periods, were comparatively insensitive to ATP removal and therefore less informative for isolating activity-dependent effects. Together, these results indicate that ATP-dependent modulation is essential for capturing the amplitude scaling, synergistic integration, and fine temporal structure of *in vivo* astrocyte calcium dynamics during active signaling epochs.

To identify which trial types contributed most to the performance decline, we compared calcium peak amplitudes predicted by the ATP-excluded and complete models (Figure S2C). For Hit, CR, and FA trials, removing ATP consistently led to smaller predicted peak amplitudes relative to both the complete model and the experimental data, indicating that ATP-dependent modulation is necessary for generating the larger calcium responses associated with locomotion and increased NE–glutamate coactivation. Miss trials showed the opposite pattern: the complete model slightly overestimated peak amplitudes, whereas the ATP-excluded model produced values comparable to the real data. For spontaneous trials, neither model captured the observed peak magnitudes, indicating that additional mechanisms—beyond those modeled here—likely contribute to these responses. Finally, the relatively modest overall changes in the global metrics, especially the lack of a significant difference in RMSE, can be explained by the predominance of spontaneous trials in the dataset, where both models performed poorly. As a result, the ATP-sensitive differences observed in Hit, CR, and FA trials had only a limited influence on the aggregate whole-trace

Figure 3. The model predicts trial-specific astrocytic calcium responses from behavior-derived molecular inputs. **(A)** Example recording showing three molecular input signals derived from run velocity, visual stimuli, and reward. NE (top) was modeled as a leaky integrator of run velocity (green curves). Glutamate (center) was modeled as impulses triggered by running and visual stimuli. DA (bottom) was modeled as impulses triggered by running and rewards. Vertical lines indicate stimulus and reward onset (salient: dashed; threshold: dotted; reward: dash-dotted). **(B)** Comparison of model-predicted (red) and real calcium signals (blue) for the example recording in (A). The two signals (bottom) show close correspondence. Vertical colored lines mark different trial types aligned at run onset. **(C-G)** Box plots showing, for each of the five performance metrics, the comparison between the complete model and the linear model across all 35 recording sessions. Each metric quantifies how well each model’s predicted calcium signal matches the real calcium signal obtained from imaging data. Lower values indicate better performance for RMSE, activity-dependent RMSE, and normalized DTW distance, whereas higher values indicate better performance for correlation and cosine similarity. Paired t-tests comparing the two models yielded the following p-values: RMSE (3.28 *×* 10^−19^), activity-dependent RMSE (6.74 *×* 10^−10^), correlation (2.98 *×* 10^−5^), cosine similarity (2.74 *×* 10^−14^), and normalized DTW distance (7.02 *×* 10^−15^). **(H)** Trial-based peak amplitudes comparison among the real calcium signals (blue), signals predicted by the linear model (green), and signals predicted by the complete model (red). The bar plot shows mean calcium peak values (*±* SEM) across the five trial types: Hit (*n* = 100, real: 0.649 *±* 0.021, linear: 0.773 *±* 0.010, complete: 0.657 *±* 0.018), CR (*n* = 135, real: 0.660 *±* 0.021, linear: 0.821 *±* 0.013, complete: 0.731 *±* 0.020), Miss (*n* = 71, real: 0.182 *±* 0.006, linear: 0.140 *±* 0.006, complete: 0.201 *±* 0.008), FA (*n* = 22, real: 0.410 *±* 0.045, linear: 0.344 *±* 0.041, complete: 0.342 *±* 0.029), and Spontaneous (*n* = 728, real: 0.270 *±* 0.005, linear: 0.353 *±* 0.009, complete: 0.281 *±* 0.004). n.s. p-value≥ 0.05; * p-value*<* 0.05; ** p-value*<* 0.01; *** p-value*<* 0.001. Error bars represent SEM. Colored dots in (C-G) denote individual recordings, and in (H) denote individual trials. measures.

### The model captures spectral features of astrocytic calcium activity

Building on the time-domain evaluations, we next examined the spectral properties of astrocytic calcium signals in the frequency domain. Time series data from all 35 recordings were concatenated, transformed using a Fast Fourier Transform (FFT), and the resulting power spectra of real calcium signals were compared with predictions from (i) the complete model, (ii) the linear model, and (iii) the fixed-ATP model (Figure S2D-G). This analysis revealed how each model distributes signal energy across frequencies.

To quantify pairwise differences between spectra, we calculated the Jensen–Shannon (JS) divergence:

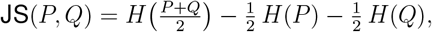

where *P*_*i*_ and *Q*_*i*_ are normalized power values (probability distributions) of real and predicted signals at frequency *i*. The Shannon entropy *H* is given by

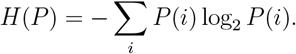

Because astrocytic calcium activity is dominated by slow dynamics in the sub-Hz range, with higher frequencies largely reflecting measurement noise, we restricted analysis to 1 ≤ Hz. The complete model reproduced the experimental spectrum with high fidelity (JS divergence = 0.0154; Figure S2E). By contrast, the linear and fixed-ATP models produced higher divergences (0.0202 and 0.0311, respectively; Figure S2F-G), indicating reduced spectral accuracy when nonlinear integration or ATP modulation is omitted.

To further assess frequency dependence, we computed cumulative JS divergence as a function of cutoff frequency (Figure S2H). Model differences were greatest at low frequencies (less than 0.1 Hz), where calcium signal power is highest, with the complete model consistently showing the lowest divergence from experimental data. Together, these results demonstrate that the complete model most effectively captures the physiologically relevant frequency content of astrocytic calcium dynamics.

### Calcium event analysis rules out ER depletion as a central mechanism of astrocytes’ refractory period

Previous studies have suggested that during large-scale calcium events, substantial efflux of calcium from the ER into the cytoplasm might deplete ER stores, with the refractory period reflecting the time required for replenishment^2,7,8,10,68^. Here, “large-scale” refers to high-amplitude transients (Δ*F/F*_0_) within or across astrocytes, often spanning broad spatial domains, such as those observed during rewarded trials (Figure S1A). Because our modeling is based on the temporal dynamics of such events, it inherently captures both temporal and spatial aspects of large-scale astrocyte activation.

To assess whether ER calcium depletion could underlie astrocytic refractory periods, we considered key biophysical constraints: the ER occupies 15–70% of the total cell volume^34–36^ and maintains calcium concentrations in the hundreds of micromolar range, whereas cytosolic calcium is typically in the hundreds of nanomolar range^33^.

We next estimated the upper bound of cytosolic calcium concentration during calcium events For each event across our 35 recordings, we quantified baseline fluorescence (*F*_0_) and Δ*F/F*_0_ Although baseline intensities varied across mice, reflecting surgical and imaging conditions, Δ*F/F*_0_ values consistently ranged from 0 to 6 (Figure S4A). Given that the reported GCaMP6f dynamic range (*F*_max_*/F*_min_ = 49.0-54.6 in vitro^69^) far exceeds these observed changes, saturation of the indicator is unlikely, even accounting for potentially lower dynamic range *in vivo*.

We then converted fluorescence signals to calcium concentrations using the Hill equation^70^:

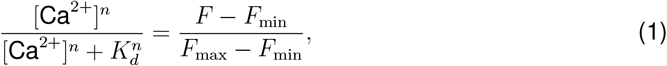

where *F* is measured fluorescence, *K*_*d*_ = 0.361-0.389 *µ*M, and *n* = 2.17-2.37^69^. For simplicity, we assumed *F*_min_ corresponds to baseline intensity and defined *F*_max_ as the largest Δ*F/F*_0_ observed across events (Figure S4A). A more detailed parameter exploration, including the effect of nonzero resting calcium (typically 50–150 nM^33,34^), is provided in Appendix S3 and Figure S4B.

Grid-search analysis revealed that most calcium elevations did not exceed 0.5 *µ*M, with an absolute upper bound of 2.618 *µ*M under extreme parameter values (*K*_*d*_ = 0.389, *n* = 2.17, [Ca^2+^]_base_ = 0.15 *µ*M). Considering that ER calcium concentrations are at least 1000-fold higher than cytosolic levels^33^ and that ER volume comprises 15–70% of the cytoplasm, cytosolic calcium would need to rise to ∼ 22.5 *µ*M to substantially deplete ER stores. Thus, the calcium events observed in our experiments are highly unlikely to deplete the ER.

We note that the Hill formulation of Eq. (1) is an approximation, and intracellular complexities—such as heterogeneous buffering, uneven indicator expression, and deviations of *in vivo* kinetics from in vitro parameters—may affect the quantitative estimates. However, these factors are unlikely to bridge the order-of-magnitude difference required to support ER depletion as a plausible mechanism.

Because ER depletion cannot explain the refractory period in astrocytic calcium signaling, we next examined alternative mechanisms, including cPKC-mediated feedback inhibition (Fig. 1E).

### Parameter fitting reveals prolonged cPKC elevation following large-scale calcium events

*in vivo* studies in the cortex and hippocampus have shown that astrocytes exhibit a refractory period lasting approximately 10–30 seconds, during which reactivation is strongly suppressed^7,8,11,12^ To investigate this phenomenon, our behavioral task incorporated a 20-second standstill phase, ensuring that Hit, Miss, CR, and FA trials were separated by ≥20 seconds, whereas Spontaneous trials involved intervals *<*20 seconds (Fig. 2B). Notably, astrocytic calcium transients in the cortex generally last between 5 and 15 seconds, with most ending within 20 seconds, thus typically exceeding brief locomotor bouts^8,9^.

To examine the relationship between inter-run interval and calcium responsiveness, we extracted ∼45-second time windows from our 35 recordings, each containing two consecutive runs (**STAR METHODS**). These windows were aligned at the onset of the second run and sorted by inter-run interval (Fig. 4A). As expected, the amplitude of the calcium response to the second run increased with longer intervals, consistent with the presence of a refractory period.

**Figure 4.**
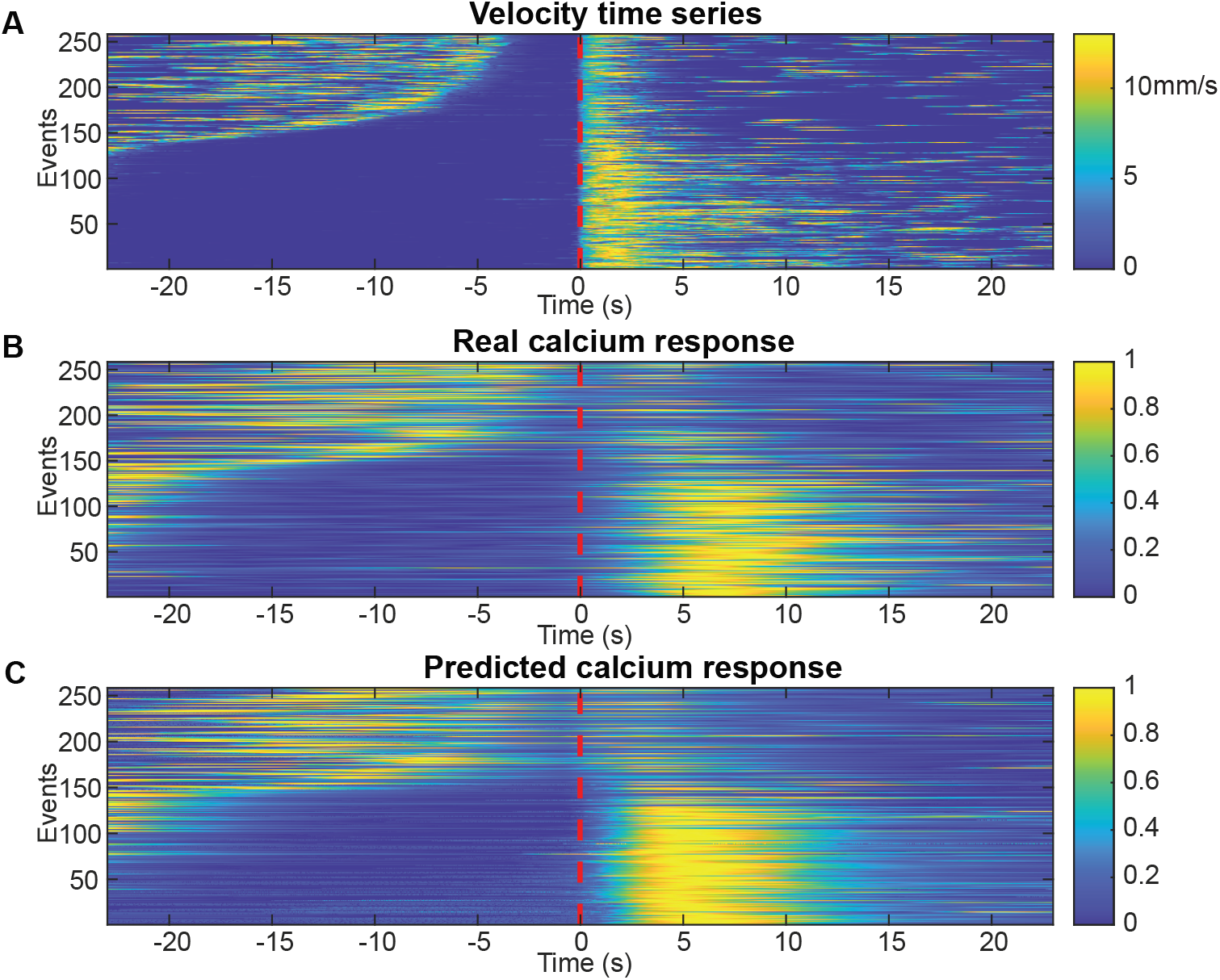
The model captures refractory periods in astrocytic calcium signals. **(A–C)** Heatmaps of run velocity (top), real calcium responses (center), and model-predicted calcium responses (bottom) for 258 inter-run intervals, aligned to the onset of the second run (red dashed line). Episodes here were selected based on the mouse’s velocity time series. Rows are sorted by inter-run interval length (ascending, top to bottom). For short intervals, real calcium responses to the second run are markedly suppressed, indicating a refractory period. The model reproduces this suppression, recapitulating the refractory dynamics observed *in vivo*.

Having ruled out ER depletion as a primary mechanism for this temporary inhibition, we considered an alternative involving negative feedback downstream of Gq-GPCR activation. Because glutamate, NE, and DA all engage Gq-coupled receptors (Fig. 1B), we focused on cPKC, which is activated by calcium and DAG, translocates to the plasma membrane, and inhibits Gq-GPCR and PLC signaling via phosphorylation^15,35,37,38^ (Fig. 1E).

In our model, cPKC dynamics were represented by a Markov chain (Appendix S1, Eq. (35)) Activation was driven by cytosolic calcium and DAG, while deactivation followed a piecewise linear time course. Specifically, once calcium exceeded a defined threshold, the deactivation rate was transiently reduced to 1% of baseline for a fixed duration *T*_cPKC_, after which it returned to normal. This period corresponds biologically to sustained cPKC activity above a functional threshold, allowing continued phosphorylation and inhibition of downstream targets.

To estimate *T*_cPKC_, we jointly optimized all model parameters within physiologically constrained ranges. For *T*_cPKC_, we set bounds of [5,100] seconds, guided by reports of a 10–20 second lag between calcium elevation and PKC activation^39^, with 100 seconds chosen as a conservative upper limit. Ranges for other parameters were determined analogously (Appendix S4).

Parameter optimization was performed using a genetic algorithm (GA)^71^ with stratified five-fold cross-validation to assess model generalization and guard against overfitting. Overfitting arises when a model learns idiosyncratic patterns and random noise specific to the training data, thereby impairing its ability to generalize and accurately predict outcomes on previously unseen test data. In our case, 35 recordings yielded 1,056 trials in total, with 98 parameters to be optimized. We applied five-fold cross-validation by stratifying the 35 recordings (7 per fold) across animals. In each round, the model was trained on four folds (28 recordings) and tested on the remaining fold, such that every recording served once as a test case (Figure S5). In each training fold, the model was fit using ∼845 trials to optimize 98 parameters, providing a favorable trial-to-parameter ratio that makes overfitting unlikely. This procedure reduces overfitting risk and ensures performance reflects generalization across animals rather than memorization. Each GA generation thus produced a distribution of *T*_cPKC_ values across folds, reflecting the evolving preference of the model during optimization. By the final GA generation, most values clustered around 20 seconds, with a minority scattered between 50–60 seconds (Fig. 5B). The strong clustering indicates a global optimum near 20 seconds, whereas scattered higher values represent local optima arising from the nonconvex optimization landscape. Together, these results suggest that cPKC remains elevated above a certain threshold for ∼20 seconds following large-scale calcium events, consistent with its proposed role in mediating the astrocytic refractory period.

**Figure 5.**
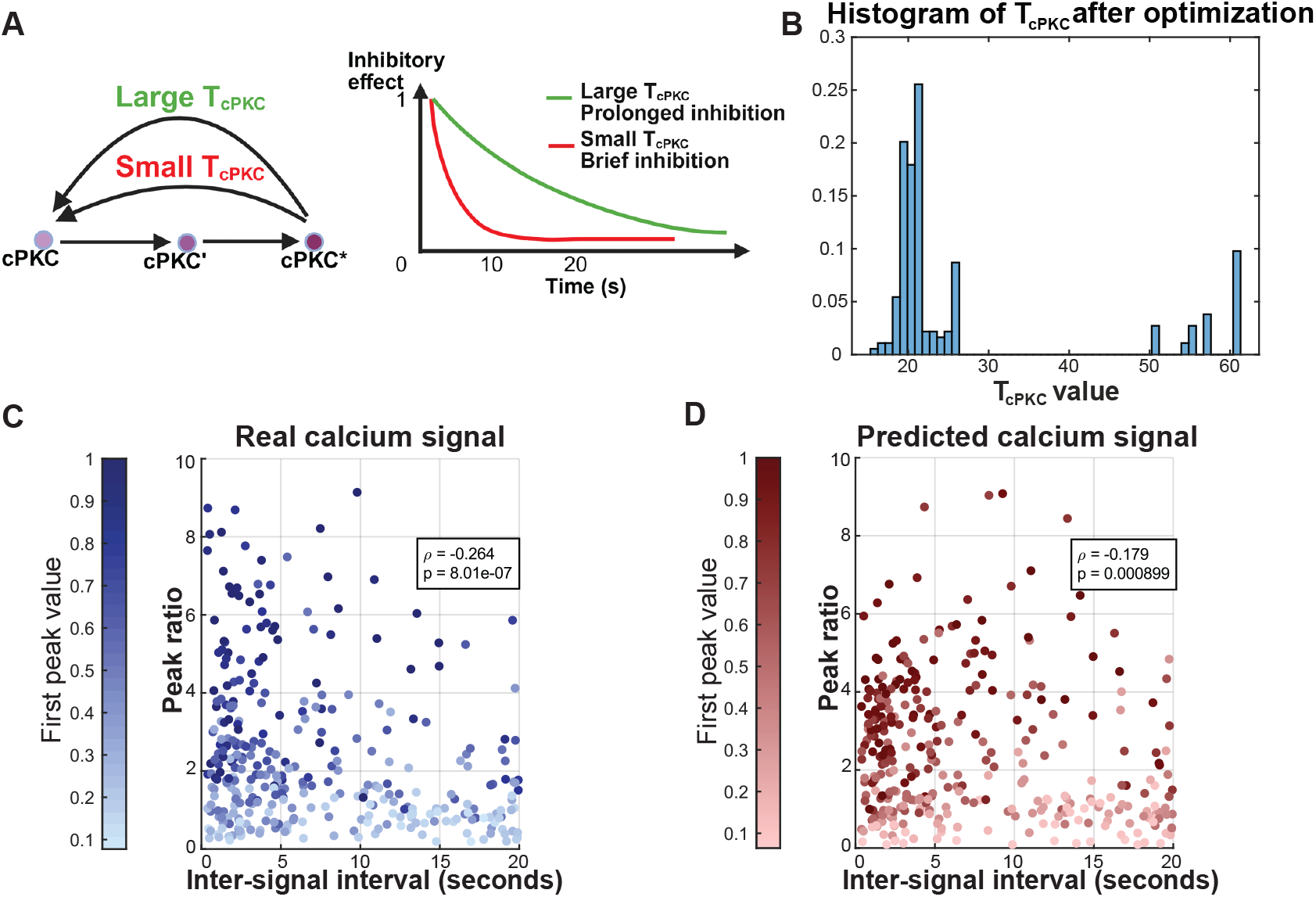
The model predicts cPKC dynamics underlying refractory suppression of calcium signals. **(A)** Schematic showing how *T*_cPKC_ influences cPKC deactivation (left) and the inhibition of Gq-GPCR–mediated calcium responses (right). Larger *T*_cPKC_ values (green) result in slower deactivation, whereas smaller values (red) produce faster deactivation. **(B)** Distribution of *T*_cPKC_ values after model optimization with a genetic algorithm (*n* = 40 population size; 15th generation). Most values clustered around 20 s, consistent with a global optimum. **(C)** Scatter plot showing the relationship between inter-signal interval and calcium peak ratio (first/second peak) in the experimental data. A significant negative correlation was observed (Spearman’s *ρ* = −0.264, p-value= 8.01 × 10^−7^; *n* = 340), indicating stronger refractory suppression at shorter intervals. The peak-pairs here were selected based on the real calcium signals. **(D)** Model-predicted relationship between inter-signal interval and calcium peak ratio, showing a similar negative correlation (Spearman’s *ρ* = −0.179, p-value= 8.99 × 10^−4^; *n* = 340). The peak-pairs here correspond to those in (C). In (C) and (D), darker dots indicate larger first-peak amplitudes. Panel (A) created with BioRender.

### A cPKC-mediated feedback mechanism explains refractory suppression of calcium responses

Using a parameter configuration with biologically plausible *T*_cPKC_ values from the final GA generation, we tested the model’s ability to reproduce astrocytic refractory dynamics during inter-trial intervals (Fig.2B). We identified pairs of calcium events that met one of two criteria: (1) the first event occurred at least 20 seconds after prior activity to ensure functional isolation, and (2) the second event followed within 20 seconds. We refer to the time gap between the pair of calcium events as the inter-signal interval, which differs from the inter-run interval shown in Fig.4 The use of inter-signal intervals reflects the fact that the refractory effect is defined by calcium events themselves; however, because the timing of such events cannot be directly controlled experimentally, this analysis provides an indirect but appropriate measure of refractory dynamics To quantify refractory strength, we calculated for each pair (340 pairs across 35 recordings) the ratio of the first peak to the second for both real and predicted calcium signals, with larger ratios indicating stronger suppression (Fig. 5C, D).

The model successfully recapitulated three key experimental features: (i) the overall magnitudes of peak ratios, (ii) the observation that larger initial peaks were often associated with greater suppression of subsequent responses, and (iii) the significant negative correlation between peak ratio and inter-event interval (real calcium: Spearman’s *ρ* = −0.264, p-value= 8.01 *×* 10^−7^; predicted calcium: *ρ* = −0.179, p-value= 8.99 *×* 10^−4^). The close match between real and predicted dynamics is further illustrated in Fig. 4A–C, showing run velocity, real calcium signals, and model predictions aligned to the onset of the second run and sorted by inter-run interval length.

To assess the specific contribution of cPKC, we repeated the analysis with cPKC removed from the signaling cascade by setting *b*_cPKC,1_ = 0 (eliminating cPKC activation). Across all five performance metrics, removing cPKC substantially degraded model accuracy, yielding higher RMSE and activity-dependent RMSE, lower correlation and cosine similarity, and increased normalized DTW distance compared to the complete model (Figure S6A-E). Moreover, the cPKC-excluded model failed to capture the negative correlation between peak ratio and inter-signal interval (*ρ* = − 0.012, p-value= 0.829) (Figure S6F), and the predicted signals lacked the characteristic suppression of the second peak (Figure S6G-I).

Together, these findings demonstrate that cPKC-mediated feedback is essential for reproducing the astrocytic refractory period and for faithfully capturing its graded suppression during inter-trial intervals.

### Linear mixed-effects analysis reveals the timescale of refractory effect decay and further disproves the ER depletion hypothesis

In addition to predicting the timescale over which cPKC remains elevated, we inferred the timescale of refractoriness decay by analyzing the relationship between the amplitude of the second calcium response (A_2_) and the inter-run interval (IRI) using a linear mixed-effects model (LME) (Figure S3 and Appendix S5), based on the data presented in Fig. 4A, B. The LME revealed that IRI (fixed effect) is a statistically significant positive predictor of A_2_, whereas the first calcium response amplitude (A_1_) and its interaction with IRI (fixed effects) are not significant. Random intercepts (random effects) were included to account for baseline differences across mice. To rule out the possibility that mice also differed in how strongly inter-run interval influenced the second calcium response, we additionally compared a random-intercept–and–slope model, but a likelihood ratio test showed no improvement in model fit (*χ*^2^(2) = 0.96, p-value = 0.62), so the simpler random-intercept model was selected. Details on the LME can be found in Appendix S5. Importantly, the lack of a significant contribution from A_1_ provides strong evidence against ER depletion as the primary mechanism of refractoriness, further supporting our previous analysis (see Appendix S3) showing that ER is unlikely to deplete during the calcium activities in our recordings. If ER calcium stores were substantially depleted by the first response, larger A_1_ amplitudes would be expected to reduce the availability of calcium for the second response, thereby exerting a strong negative effect on A_2_. The fact that A_1_ and its interaction with inter-run interval were not significant predictors in the LME instead supports the interpretation that the refractory effect arises from regulatory feedback mechanisms, such as cPKC activity, rather than depletion of ER calcium stores.

To further interpret these results, we considered two characteristic recovery points: the time required for the refractory effect to fully decay and the time for half decay. The LME estimated that the refractory effect fully decays when IRI is approximately 63 seconds and half degraded at about 32 seconds (Figure S3A), consistent with the results in literature^8,9^. A detailed breakdown of the sequential processes underlying these timescales is provided in Appendix S5, where we further estimated that cPKC requires roughly 7 seconds of partial degradation to reduce its inhibitory effect on calcium by half.

## DISCUSSION

Astrocytic calcium dynamics are shaped by diverse neuromodulatory and synaptic inputs, yet their temporal integration is constrained by a refractory period that has remained mechanistically unresolved. Here, we developed a compartmental model linking behavior-derived glutamate, NE, and DA inputs to intracellular calcium signaling, and identified a cPKC–mediated feedback as a parsimonious mechanism underlying refractoriness. Notably, we arrived at this mechanism only after rigorously excluding ER depletion as a viable explanation using two independent analyses of *in vivo* data. The model, validated against *in vivo* calcium imaging data from behaving mice (Fig. 2), reproduces trial-dependent calcium dynamics across multiple metrics (Fig. 3) and explains the graded suppression of responses observed at short inter-event intervals (Figs. 4, 5). Our findings indicate that the refractory period is not a passive consequence of ER depletion but an actively enforced regulatory state. By constraining when astrocytes re-enter an excitable mode, the refractory period fundamentally shapes their ability to modulate synapses and circuit activities.

Our results demonstrate that nonlinear interactions between signaling pathways are crucial for accurately predicting astrocytic calcium dynamics. Ablating nonlinear terms or fixing ATP at constant level reduced performance, underscoring the role of ATP in enabling the synergistic integration of NE and glutamate. These findings support experimental evidence that NE gates glutamatergic and dopaminergic responses^4^, and suggest that metabolic state sets the dynamic range within which astrocytes integrate synaptic and neuromodulatory inputs^72^. However, direct verification of the modeled NE–DA gating effect cannot be achieved with the current experimental dataset, as this would require isolating dopaminergic inputs and comparing calcium responses in the presence and absence of concurrent NE stimulation.

ER depletion has been proposed as a possible explanation for astrocytic refractoriness^7,8,10,11^ However, quantitative analyses of event amplitudes and ER calcium capacity suggest that this mechanism is unlikely under our conditions. Moreover, our linear mixed-effects model indicated that, after accounting for the inter-run interval, the first calcium amplitude was not a significant predictor of the second. This further argues against ER depletion as the underlying cause of refractoriness. The two independent data-mining approaches jointly indicated that an alternative mechanism must contribute to the calcium refractory period. To address this, we proposed the cPKC-mediated negative feedback on Gq-GPCR and PLC as a primary cause, whose ablation rendered the model unable to capture the refractory period. We used the model to predict that cPKC remains elevated for ∼20 seconds after large calcium events, while the linear mixed-effects model indicated that the refractory effect is half-degraded at an inter-run interval of ∼32 seconds Within this interval, we further estimated that cPKC requires ∼7 seconds of partial degradation after activation to reduce its inhibitory effect on calcium by half, thereby forming a feedback gate at the Gq/PLC level. This finding captures key experimental features, including stronger suppression following large initial peaks and recovery with longer inter-event intervals. Such gating may help prevent excessive gliotransmitter release, maintain stable circuit activity, and ensure that astrocytes preferentially respond to the most behaviorally relevant signals.

Future optical PKC reporters are expected to reveal ∼10–30 s elevations in activity following large calcium events, with decay kinetics (including a half-time of ∼7 s) corresponding to the recovery of astrocytic responsiveness. The model further predicts that pharmacological or genetic inhibition of PKC will shorten the refractory period and enhance calcium responses to closely spaced inputs, whereas PKC activation will extend suppression. In addition, our framework highlights the dual role of ATP. While our model formalizes ATP primarily at the level of its production, astrocytic ATP concentrations are normally maintained in the millimolar range^50^, where ATP exerts an inhibitory influence on IP_3_R2 gating. Under disease conditions, impaired ATP production can lower cytosolic ATP into the sub-millimolar range^50^, where ATP instead facilitates IP_3_R2 activity. This suggests that ATP depletion, although detrimental to cellular energetics, may paradoxically enhance receptor-mediated calcium release and alter nonlinear input integration in astrocytes.

Future validation will benefit from genetically encoded sensors for NE^73^, DA^55^, and ATP^74^, which will provide direct measurements of the molecular inputs currently inferred from behavior. The observed trial-type biases also suggest that additional factors, including acetylcholine, serotonin, or unmeasured behaviors, may contribute to spontaneous and false-alarm responses Moreover, several modulatory pathways not yet included in our model, such as P2X receptor–mediated calcium entry, receptor trafficking, and longer-timescale NE–DA interactions^4,41^, likely play important roles. Our mean-field framework also cannot capture spatial microdomain heterogeneity or stochastic channel gating. Ultimately, next-generation indicators for transmitter release will enable ground-truth tests of how the spatial and temporal features of astrocytic excitation shape neural circuit activity.

Recent advances in neural surrogates and operator-learning frameworks present exciting new opportunities to accelerate or augment mechanistic models of intracellular signaling. Our simulation framework generates high-resolution flux and state trajectories that could serve as rich training data for neural operator or state-space surrogate models, enabling real-time prediction of astrocytic calcium dynamics from diverse neuromodulatory inputs. Such AI-augmented extensions, which couple the interpretability and mechanistic insight of biophysical models with the computational efficiency of data-driven approaches, represent a promising direction for future work.

In summary, by linking behaviorally derived molecular inputs to intracellular signaling and validating model predictions against *in vivo* data, this study establishes a behavior-grounded “virtual astrocyte” framework. Extending this approach to incorporate spatially resolved signaling domains, multicellular network-level interactions, and disease-relevant perturbations will provide deeper insight into how astrocytes integrate diverse molecular cues to modulate neural circuit dynamics. More broadly, the paradigm introduced here highlights the critical roles of nonlinear feedback, metabolic coupling, and neuromodulatory context in shaping glial responsiveness to behaviorally relevant activity, and offers a scalable modeling strategy for bridging molecular mechanisms with emergent systems-level phenomena.

## RESOURCE AVAILABILITY

### Lead contact

For further information and requests for resources or code, contact Axel Nimmerjahn or Guoqiang Yu.

### Materials availability

This study did not generate new, unique reagents.

### Data and code availability

The full implementation of the model, including simulation scripts and the processed data files, is available at [GitHub repository](https://github.com/jameschengpeng/Calcium_model_explaining_refractory.git).

## ACKNOWLEDGMENTS

J.C.P. gratefully acknowledges the invaluable assistance of Boyu Lyu in quantifying the mitochondrial-to-cytoplasmic volume ratios, Xuelong Mi for expert guidance in operating the AQuA2 software, Wei Zheng for modeling suggestions, and Jiayi Zhao for guidance on building the linear mixed-effect model. J.C.P., G.Y., and A.N. gratefully acknowledge Shaul Druckmann, Nathan Smith, and D.P. Mohapatra for their valuable input throughout the development of this project. This work was primarily supported by the National Institutes of Health (NIH) grant U19 NS123719 (to L.T., A.N., and G.Y.). It was partially supported by NIH grant R01 MH110504 (to G.Y.) and T32 GM007198 (to M.K.D.); the Edwards-Yeckel Research Foundation and NOMIS Neuroimmunology Initiative (to A.N.), and a Dan and Martina Lewis Biophotonics Fellowship (to M.K.D.). The content is solely the authors’ responsibility and does not necessarily represent the official views of the NIH or other funding agencies.

## AUTHOR CONTRIBUTIONS

J.C.P. designed and implemented the calcium signaling model, performed all computational analyses, and wrote the manuscript. K.M. conducted the *in vivo* experiments and provided the calcium imaging data used for model evaluation. M.K.D., N.N., Yizhi Wang, and D.D. reviewed the manuscript and provided critical feedback. Yue Wang contributed suggestions specifically related to parameter optimization strategies. L.T. provided guidance on deriving physiological bounds for calcium concentrations based on fluorescence imaging data. A.N. and G.Y. co-led the project and provided supervision throughout.

## DECLARATION OF GENERATIVE AI AND AI-ASSISTED TECHNOLOGIES

During the preparation of this work, the author(s) used ChatGPT in order to polish the writing. After using this tool or service, the author(s) reviewed and edited the content as needed and take(s) full responsibility for the content of the publication.

## SUPPLEMENTAL INFORMATION

### Supplementary Figures

**Figure S1.**
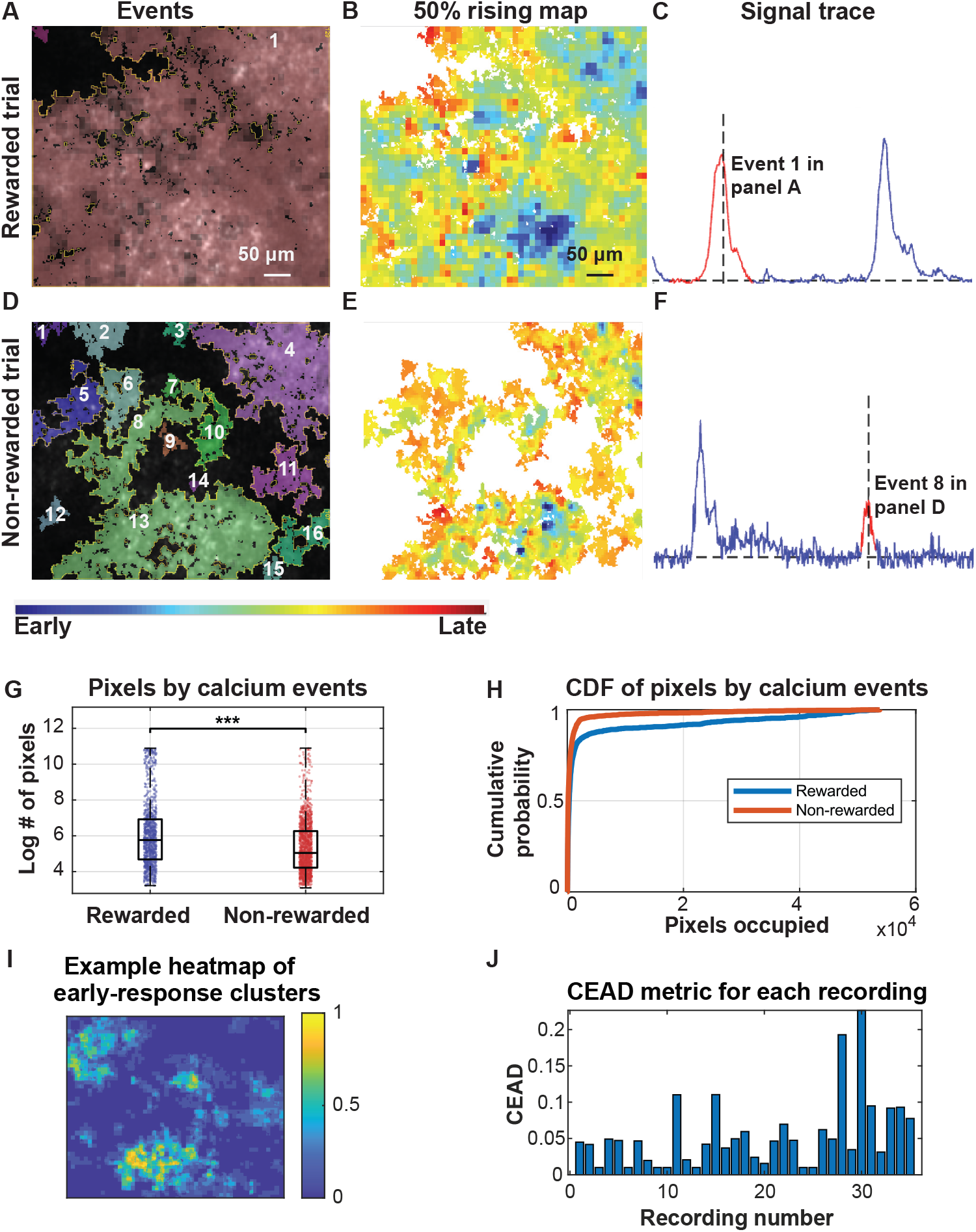
AQuA2 analysis reveals trial-type-dependent calcium dynamics and spatial clustering of early response regions, related to Figure 2. **(A)** A large calcium event was detected in a rewarded trial. The event index is shown within the segmented region. **(B)** The 50% rising map of the pixels occupied by the event shown in (A), indicating the time at which each pixel reaches 50% of its amplitude. Blue represents earlier timings, while red represents later timings **(C)** The signal trace corresponding to the region occupied by the event in (A). **(D)** Multiple smaller calcium events observed in a non-rewarded trial, each outlined with a yellow boundary. Event indices are labeled within each segmented region. **(E)** The 50% rising map of the pixels in (D), with color indicating the relative timing of pixel activation. **(F)** The signal trace for one specific event, number 112, from (E). **(G)** Log-transformed box plot showing the number of pixels that are occupied by calcium events associated with rewarded trials (left) and non-rewarded trials (right) Among the 35 recordings, there are 201 rewarded trials, containing 1134 calcium events (detected by AQuA2), and 344 non-rewarded trials, with 1447 calcium events. The unpaired two-sample t-test has a 95% confidence interval [2240, 3473] with p-value 5.203 × 10^−13^, which indicates that the calcium events in rewarded trials occupy a significantly larger number of pixels than those in non-rewarded trials. **(H)** The empirical cumulative distribution function of pixels occupied by calcium events in rewarded and non-rewarded trials. Since the curve for rewarded trials is beneath the one for non-rewarded trials, the rewarded ones have a heavier tail, i.e., more calcium events that occupy a large spatial domain. **(I)** An example heatmap after normalization, showing the frequency of early reactions for each pixel across calcium events in a single recording. Each pixel value represents how often it reacted early during calcium activities, revealing clustered regions of consistent early responses. This clustering suggests that specific areas within the FOV consistently respond early, potentially indicating proximity to neuronal release sites or biologically significant regions. **(J)** A bar plot displaying the CEAD scores for all 35 calcium recordings. The CEAD metric was calculated by identifying pixels in the heatmap that consistently reacted early (exceeding a threshold of 50% of the maximum early-response value), clustering them using connected component analysis, and summing the total area of these clusters. The CEAD score for each recording is defined as: 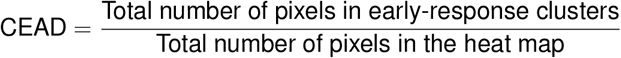 A higher CEAD score indicates more widespread early activation throughout the field of view (FOV), while a lower CEAD score reflects more localized early-response regions. Since all CEAD scores are below 0.25, with most falling below 0.1, this suggests that early reaction regions are highly localized. This localization supports the rationale for selecting the largest early reaction region to compute the reference calcium signal, as it consistently reacts rapidly following synaptic or neuromodulatory signals. n.s. indicates p-value ≥ 0.05; * Indicates p-value *<* 0.05; ** indicates p-value *<* 0.01; *** indicates p-value *<* 0.001.

**Figure S2.**
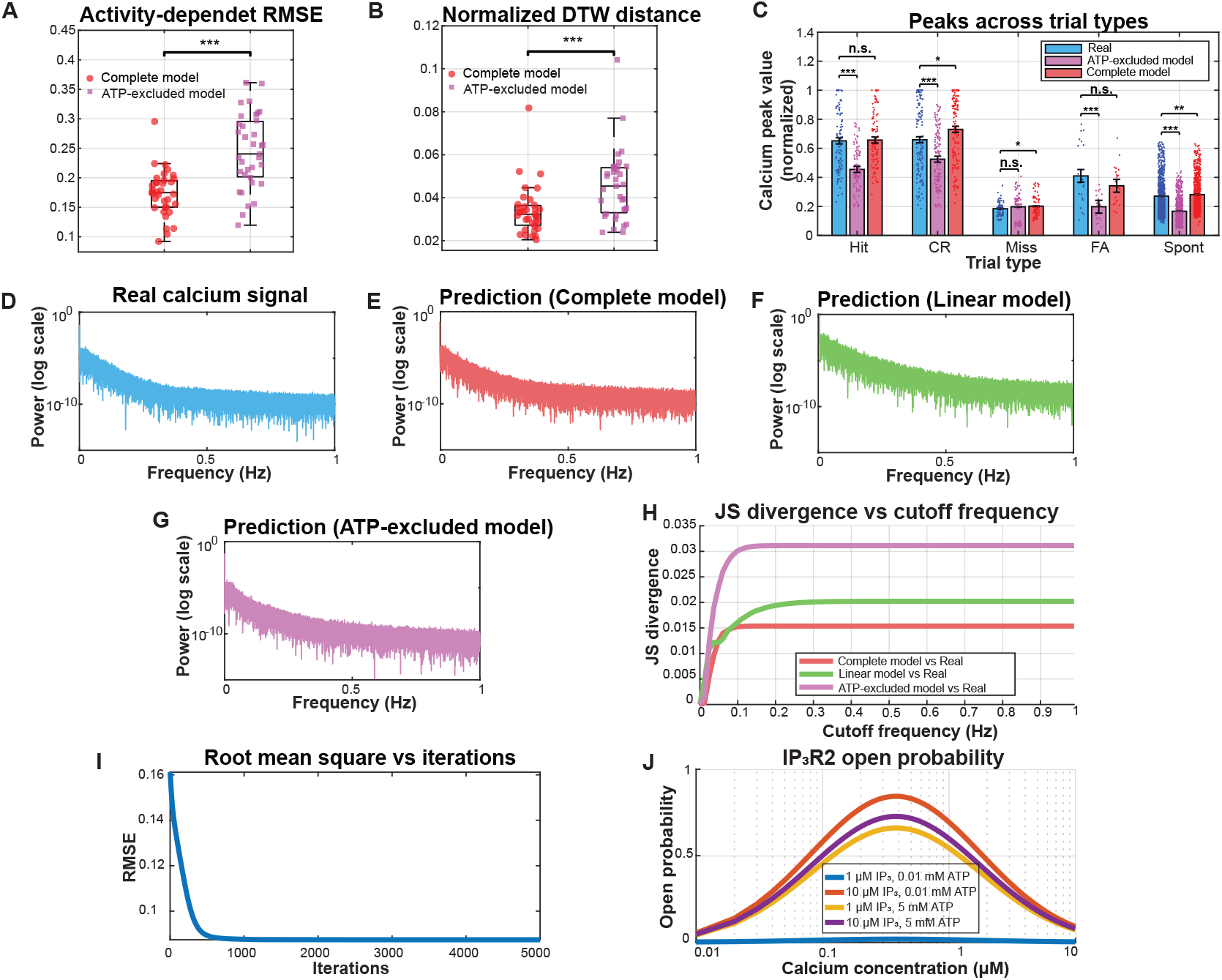
The complete calcium model outperforms linear and ATP-excluded models and captures frequency features, related to Figure 3. **(A-B)** Box plots comparing the performance of the complete model and the ATP-excluded model across all 35 recording sessions using two activity-sensitive metrics: (A) activity-dependent RMSE and (B) normalized DTW distance. Paired t-tests revealed significantly lower errors for the complete model relative to the ATP-excluded model for both metrics (activity-dependent RMSE, *p* = 7.20 × 10^−9^; normalized DTW distance, *p* = 3.88 × 10^−7^). **(C)** Trial-based peak value comparison among the real calcium signals (blue), signals predicted by ATP-excluded model (purple), and signals predicted by the complete model (red). Removing ATP from the model reduces the synergistic effect between NE and glutamate, yielding significantly smaller peak values on Hit and CR trials. The bar plot shows mean calcium peak values (± SEM) across five trial types: Hit (*n* = 100, real: 0.649 *±* 0.021, ATP-excluded: 0.455 *±* 0.014, complete: 0.657 ± 0.018), CR (*n* = 135, real: 0.660 *±* 0.021, ATP-excluded: 0.525 *±* 0.017, complete: 0.731 *±* 0.020), Miss (*n* = 71, real: 0.182 *±* 0.006, ATP-excluded: 0.197 *±* 0.011, complete: 0.201 *±* 0.008), FA (*n* = 22, real: 0.410 *±* 0.045, ATP-excluded: 0.198 *±* 0.018, complete: 0.342 *±* 0.029), and Spontaneous (*n* = 728, real: 0.270 *±* 0.005, ATP-excluded: 0.167 *±* 0.003, complete: 0.281 *±* 0.004). **(D)** Log-scaled power spectral density of the real calcium response, restricted to the physiologically relevant frequency range (≤1 Hz) **(E)** Log-scaled power spectral density of the prediction by the complete model, restricted to the physiologically relevant frequency range (≤ 1 Hz), with a Jensen-Shannon divergence to the real signal of 0.0045. **(F)** Log-scaled power spectral density of the prediction by the linear model, restricted to the physiologically relevant frequency range (≤1 Hz), with a Jensen-Shannon divergence to the real signal of 0.0068. **(G)** Log-scaled power spectral density of the prediction by the ATP-excluded model, restricted to the physiologically relevant frequency range (≤1 Hz), with a Jensen-Shannon divergence to the real signal of 0.0049. **(H)** Cumulative Jensen–Shannon (JS) divergence between the real calcium response and three model predictions as a function of cutoff frequency *f*_max_ (linear power spectral density). The three curves correspond to the complete model (red), the linear model (green), and the ATP-excluded model (pink). Frequencies up to 1 Hz are considered, and the JS divergence at each *f*_max_ is computed using only the power spectral density components within the specified frequency range. **(I)** Propagation of root mean square error (RMSE) for parameter fitting in the joint-effect model on IP_3_R2. The RMSE is calculated by comparing the model-fitted open probabilities of IP_3_R2 against real open probabilities from single-channel data under different concentration combinations of IP_3_, ATP, and calcium. The model parameters are optimized by the gradient descent method. Derivations can be found in Appendix S2. **(J)** The open probability of IP_3_R2 computed by the join-effect model based on the fitted parameters. n.s. p-value *>*0.05; * p-value *<* 0.05; ** p-value *<* 0.01; *** p-value *<* 0.001. All error bars represent SEM.

**Figure S3.**
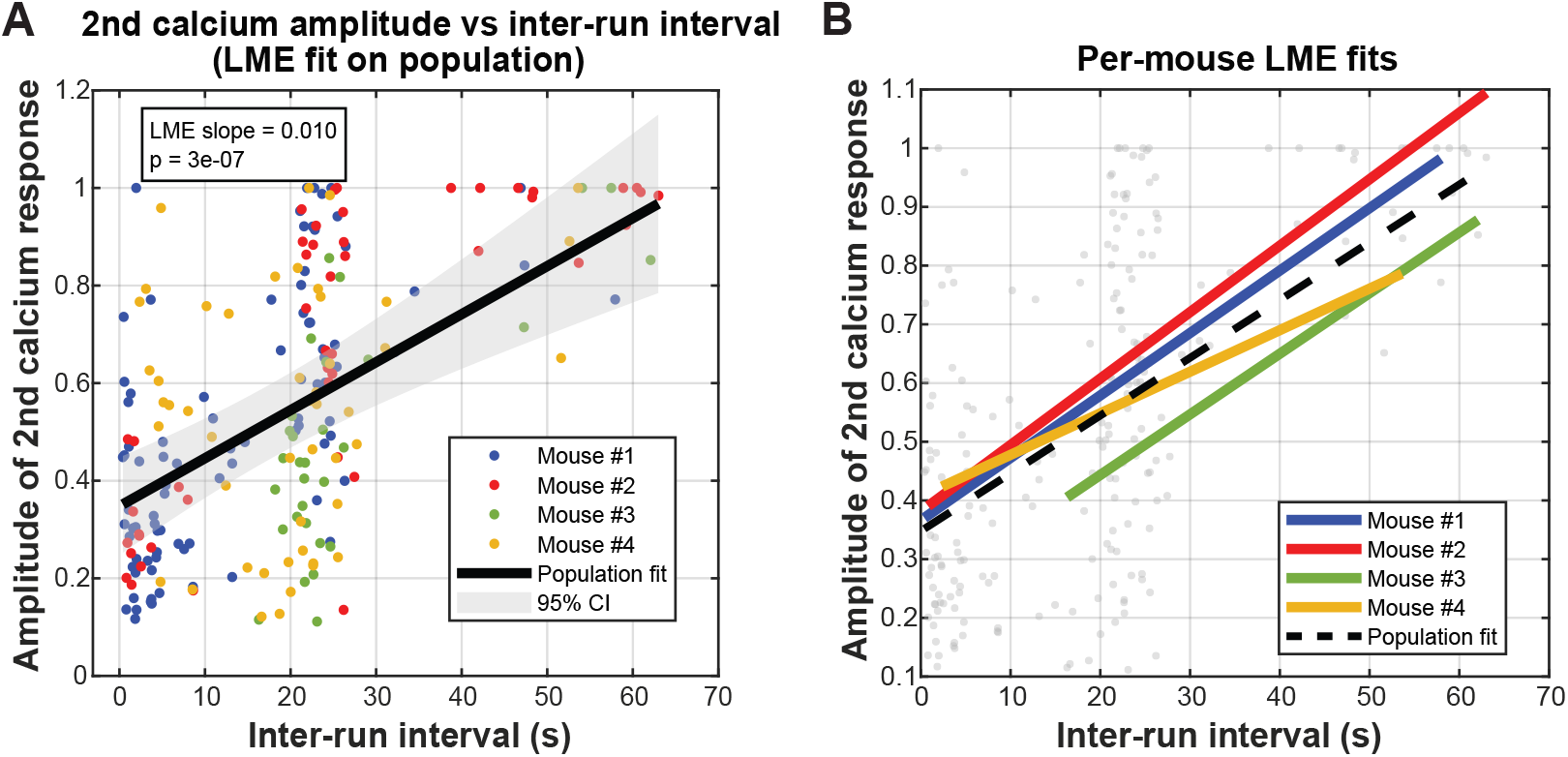
Statistical analysis reveals that the amplitude of astrocytes’ calcium response positively correlates with inter-run interval, related to Figure 4. **(A)** Linear mixed-effects model (LME) fit on the population shows that longer inter-run intervals (fixed effects) are associated with larger amplitudes in the second calcium response (slope = 0.010, p-value = 4.5 × 10^−7^), whereas the amplitude of the first calcium response and its interaction with inter-run interval (fixed effects) are not significant predictors, with p-values 0.28 and 0.34 respectively. The p-values stated here are from a simple regression on the population, whereas the full LME results, which account for mouse-to-mouse variability, are reported in Appendix S5. Random intercepts (random effects) are incorporated to account for baseline differences across mice. Shaded area denotes 95% confidence interval. The fitted line indicates a baseline amplitude of approximately 0.34 at an inter-run interval of 0 second, with the refractory effect completely degraded by around 63 seconds when the second-event amplitude reaches 1. By interpolation, the refractory effect decays to half of its magnitude at around 32 seconds. **(B)** Per-mouse LME fits demonstrate that this positive relationship is consistent across individual animals, with variation in intercepts, demonstrating the necessity of including random intercepts for each mouse. See Appendix S5 for details on the LME.

**Figure S4.**
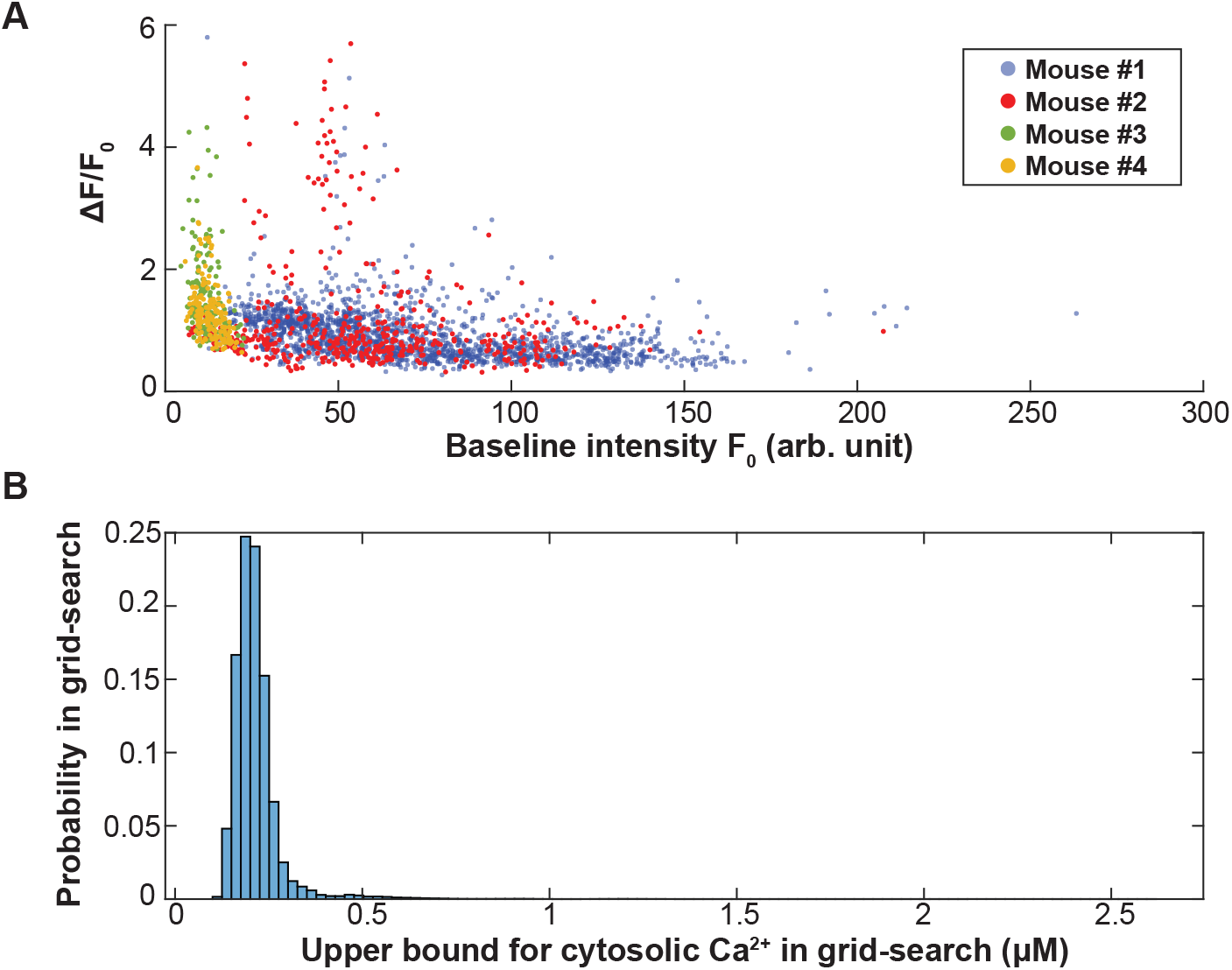
ER stores are not depleted during behavior-evoked calcium transients, related to Figure 5. **(A)** Relationship between baseline intensity and Δ*F/F*_0_ for calcium events detected by AQuA2 on the 35 recordings from the four mice. Mouse #1 (with 14 recordings and *n* = 1791 calcium events) and #2 (with 11 recordings and *n* = 490 calcium events) exhibit a broader range of baseline intensities, yet Δ*F/F*_0_ remains constrained below 6, indicating that GCaMP6f does not reach saturation even at higher signal levels. Mouse #3 (with five recordings and *n* = 151 calcium events) and #4 (with five recordings and *n* = 158 calcium events) have lower signal-to-noise ratios Despite lower baseline intensities, Δ*F/F*_0_ remains within the same bound, further supporting that GCaMP6f is not saturated across different signal levels. Overall, the maximum Δ*F/F*_0_ is 5.8 while the minimum *F*_0_ is 4.5. **(B)** Histogram of the upper bound of cytosolic calcium after conducting a grid-search on the parameter space (size *n* = 6699 after discretization) of GCaMP6f and the range of baseline calcium levels. See Appendix S3 for more details on this analysis.

**Figure S5.**
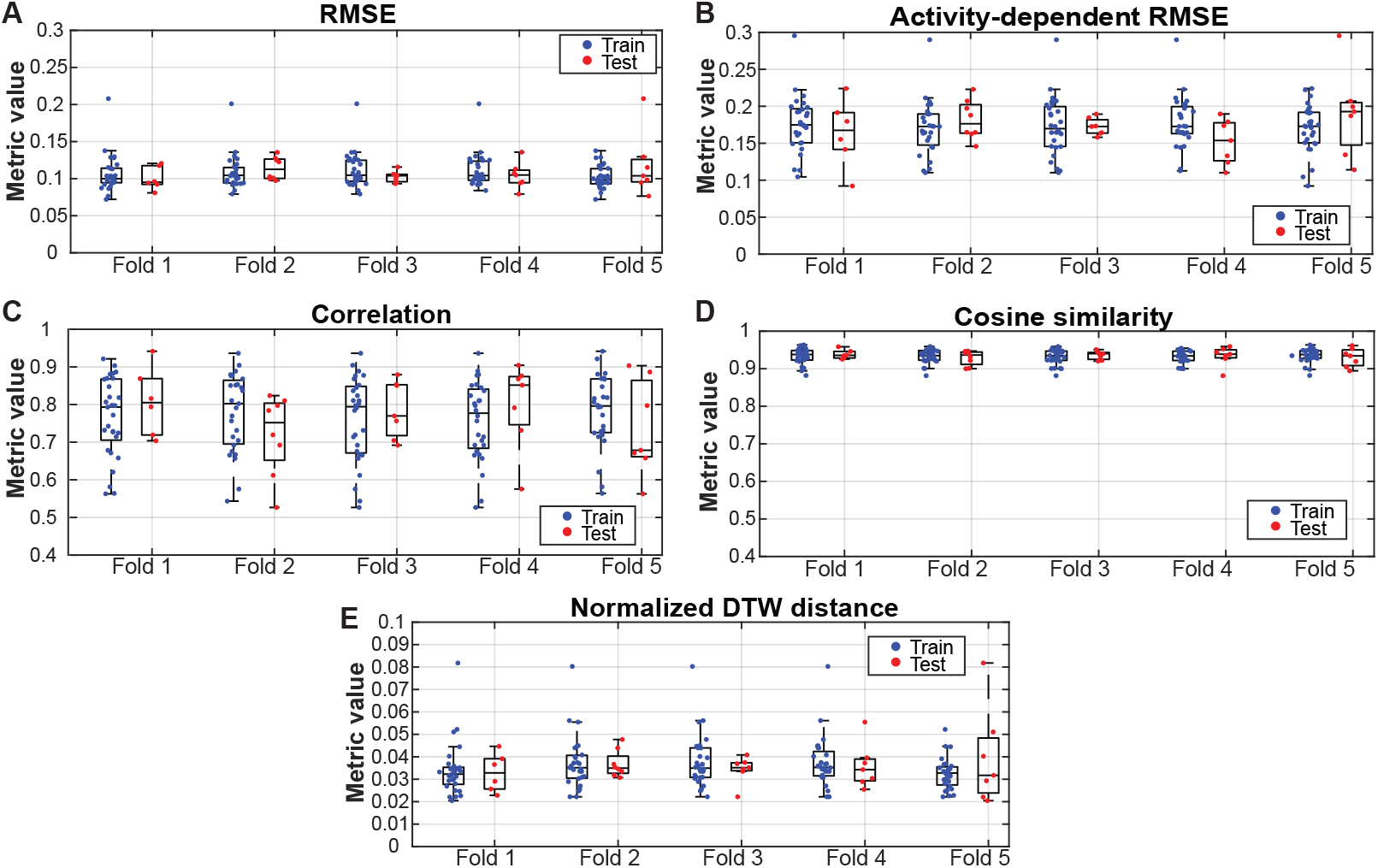
Cross-validation confirms model robustness and mitigates overfitting, related to Figure 5. To guard against overfitting and to evaluate the model’s generalizability and robustness, we performed a five-fold stratified cross-validation, where each fold preserves the proportion of recordings originating from each mouse. In each round, model parameters were optimized using recordings from four folds (training set, *n* = 28) and evaluated on the remaining fold (test set, *n* = 7). Across all 35 recordings, the dataset comprised 1,056 trials in total, while the model contained 98 parameters; in each fold, the training set contained roughly 845 trials, yielding a favorable data-to-parameter ratio that makes overfitting unlikely. For each fold, the final selected parameter configuration was chosen from the Pareto front obtained in the last generation of the genetic algorithm, specifically the configuration with the smallest root-mean-squared error (RMSE) on the training set. **(A–E)** show the model’s performance in terms of (A) RMSE, (B) activity-dependent RMSE, (C) correlation, (D) cosine similarity, and (E) normalized dynamic time warping (DTW) distance. RMSE measures the average pointwise deviation between the predicted and recorded calcium traces across the entire signal, with smaller values indicating higher overall accuracy. Activity-dependent RMSE quantifies the prediction error specifically during calcium events, excluding quiescent periods and therefore capturing performance on physiologically meaningful transients. Correlation evaluates temporal synchrony between the predicted and recorded signals; values closer to 1 indicate tighter alignment in timing. Cosine similarity treats each calcium trace as a vector and measures the cosine of the angle between them, providing a scale-independent assessment of waveform similarity; values near 1 indicate strong agreement Normalized DTW distance measures the temporal similarity between predicted and real calcium traces by computing the DTW alignment cost and dividing by trace length; values below 0.05 indicate an almost identical match, 0.05–0.10 a good match, 0.10–0.20 a moderate mismatch, and values above 0.20 reflect poor alignment. Each point represents one recording, with blue and red indicating training and test performance, respectively, and box plots summarizing the distributions across folds. The consistent generalization across all folds confirms that the model does not overfit and performs robustly on unseen recordings.

**Figure S6.**
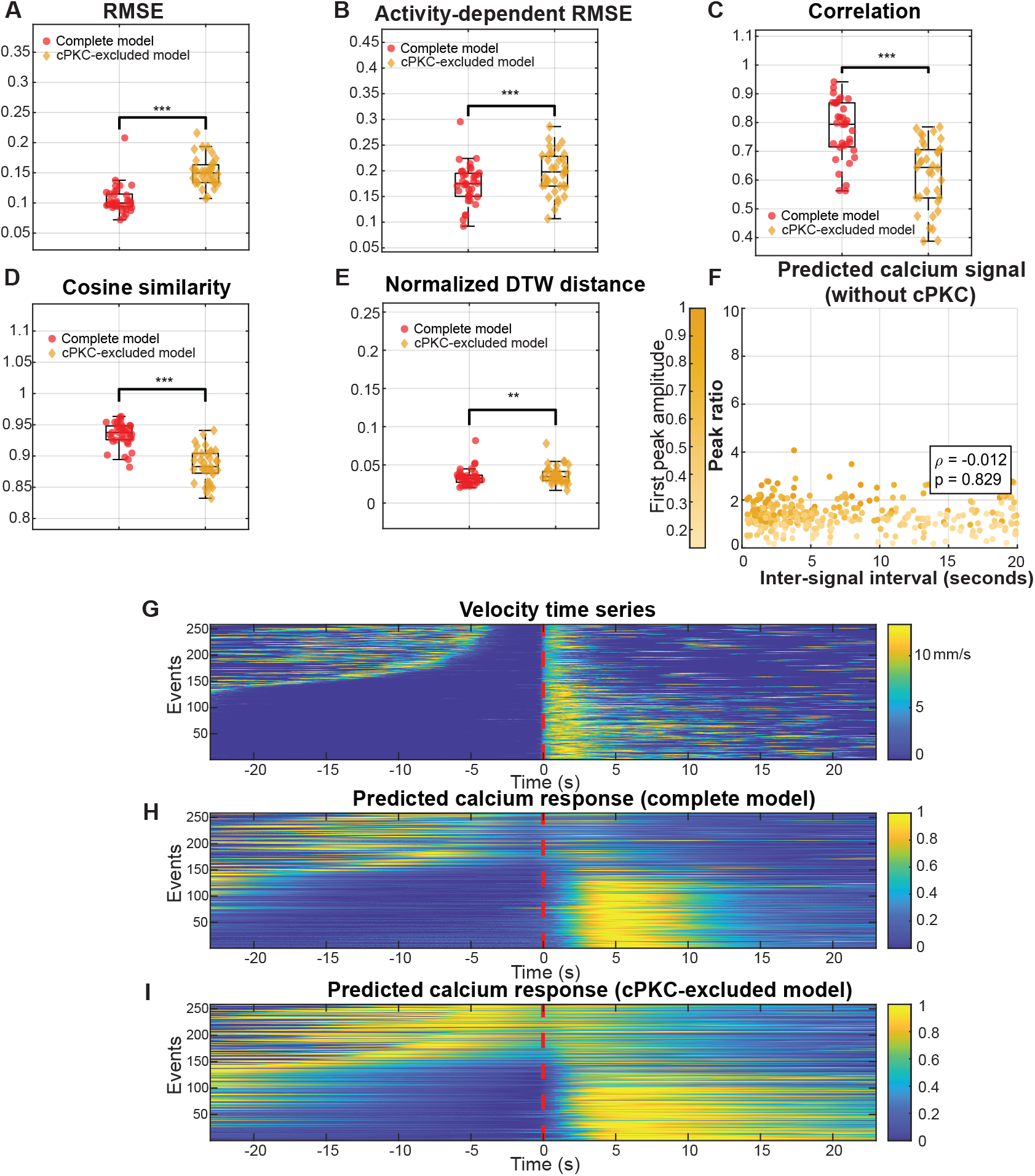
Removing cPKC reduces prediction accuracy and abolishes the model’s capability to capture the refractory period, related to Figures 4 and 5. To evaluate the significance of incorporating cPKC into the model, we removed it by setting *b*_cPKC,1_ = 0, where *b*_cPKC,1_ is the coefficient governing cPKC activation, thereby fixing cPKC at a constant value of zero. **(A-E)** Box plots showing, for each of the five performance metrics, the performance of the complete model and the cPKC-excluded model across all 35 recording sessions. Across the five performance metrics, lower values indicate better performance for RMSE, activity-dependent RMSE, and normalized DTW distance, whereas higher values indicate better performance for correlation and cosine similarity. Paired t-tests comparing the complete and ATP-excluded models yielded the following p-values: RMSE (2.80 × 10^−13^), activity-dependent RMSE (2.53 × 10^−5^), correlation (8.31 × 10^−17^), cosine similarity (4.45 × 10^−15^), and normalized DTW distance (4.75 × 10^−3^) Including cPKC results in significantly better performance across all metrics, highlighting the importance of cPKC-mediated regulation in modeling astrocytic calcium signaling. Removing cPKC impairs the model’s ability to capture the underlying dynamics, leading to reduced predictive performance. **(F)** After removing cPKC, the model shows no significant negative correlation between the inter-signal interval and calcium peak ratio (first peak value divided by the second), indicating that it fails to capture the refractory period. In the scatter plot, darker colors correspond to larger first peak values. **(G)** Heatmap of the mouse’s run velocity (mm/s), identical to Fig. 4A There are 458 traces in total. Traces were aligned at the second run-onset. **(H)** Heatmap of the predicted calcium response from the complete model, identical to Fig. 4C. Traces were aligned at the second run-onset. **(I)** Corresponding predicted normalized calcium response without cPKC, showing a much weaker refractory effect compared with the complete model in Fig. 4C. Traces were aligned at the second run-onset. n.s. p-value ≥ 0.05; * p-value *<* 0.05; ** p-value *<* 0.01; *** p-value *<* 0.001. All error bars represent SEM.

## Appendix S1

### Calcium mass conservation across compartments

In our modeling of calcium signals, we consider four compartments: ER, cytoplasm, mitochondria, and extracellular space. Given that the calcium concentration in the extracellular space is on the order of a few millimolars while that in cytoplasm is on the order of tens of nanomolars, we can reasonably assume it to be effectively infinite compared to the cytosolic calcium concentration^34^ The time derivative of cytosolic calcium in astrocytes is modeled by Eq. (2), which represents a linear combination of several key fluxes: calcium efflux via IP_3_R2 (*J*_ER_), calcium influx via the sarcoplasmic/endoplasmic reticulum calcium-ATPase (SERCA) pump (*J*_SERCA_), calcium leakage through the ER membrane (*J*_leak_), calcium influx and efflux across the plasma membrane (*J*_in_ and *J*_out_), calcium efflux from mitochondria through the mPTP 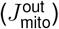 ^75^, and calcium influx from the cytoplasm into mitochondria across the mitochondrial membrane 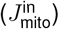.

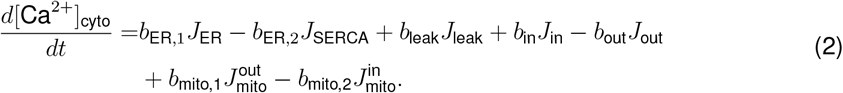

Using the fluxes in Eq. (2), we can define the time-derivative of calcium in ER and mitochondria in a similar way, as shown in Eqs. (3), (4). The constants *γ*_1_, *γ*_2_ are the volume ratios for cytoplasm-to-ER and cytoplasm-to-mitochondria. A common value for *γ*_1_ is around 5^36^. As for *γ*_2_, according to our analysis on electron microscopy data for astrocytes using the VSOT method^76^, it ranges from 16 to 20. We adopted the value 20 for *γ*_2_ in this study.

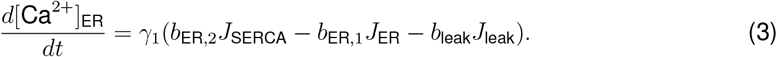

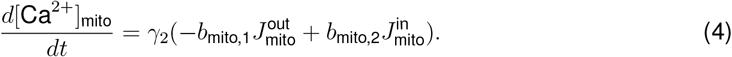

Eqs. (2), (3), (4) provide a simple yet efficient way to model the calcium dynamics in different compartments using the law of conservation of mass. Generally, the calcium concentration in astrocytes’ cytoplasm ranges between 50-150 nM^34^, while in the ER, it is around 50 *µ*M^33^. In mitochondria, calcium concentration varies significantly based on the cell’s activation state. Under resting conditions, the calcium concentration in the mitochondrial matrix is approximately the same as in the cytoplasm. However, upon activation, it can increase by 10-20 fold^51^. Thus, the mitochondria serve as a calcium buffer upon stimulation.

Next, we detail the individual calcium fluxes grouped by source: ER, mitochondria, and plasma membrane.

#### Calcium flows related to the endoplasmic reticulum

Calcium can be released from the ER via various channels, including IP_3_R1, IP_3_R2, and ryanodine receptors (RyR). Specifically in mouse brain astrocytes, the IP_3_R2 plays a major role, while the gene expression levels of IP_3_R1 and RyR are negligible according to the GeneCards database^57^ and the Brain RNA-Seq database^40^. The dynamics of IP_3_R2 have been extensively modeled to understand cellular calcium release. While early models focused on deterministic descriptions of average calcium fluxes, later stochastic models, including Markov models, provided more detailed representations However, most existing models emphasize single-channel behavior, which is not aligned with our focus on the overall effect of IP_3_R activity on calcium signaling at a macroscopic level. Additionally, these models are often parameterized for specific concentration ranges of calcium, IP_3_, and ATP, limiting their applicability across different physiological conditions. To address these limitations, we developed a model that accommodates a wider range of chemical concentrations, providing a more comprehensive representation of IP_3_R dynamics.

Our model for calcium efflux from the ER considers the combined effects of cytosolic calcium, ATP, IP_3_, ROS, and PKA. This approach incorporates more factors than previous models, allowing for a versatile and adaptable representation of IP_3_R activity. We used data from the literature^45^ to parameterize the joint effects of cytosolic calcium, ATP, and IP_3_ on IP_3_R2, and modeled the influence of cytosolic ROS and PKA using second-order Hill functions. The calcium efflux from the ER is modeled by Eq. (5),

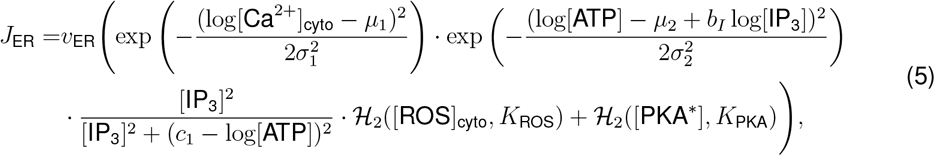

where *v*_ER_ is the maximum efflux rate. A more detailed elaboration of this model can be found in Appendix S2.

The modeling of calcium influx into the ER via the calcium-ATPase pump (SERCA) is much simpler than that of efflux. We refer to literature^77^, and used Eq. (6):

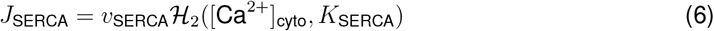

Besides the IP_3_R and SERCA, there is also calcium leakage from the ER to the cytoplasm via the ER membrane. We assume that the leakage is dependent only on the concentration gradient between the ER and cytoplasm, which yields Eq. (7):

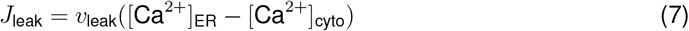

#### Calcium flows related to mitochondria

In this section, we introduce the modeling approach for calcium dynamics related to mitochondria. The efflux occurs through the mPTP, which is activated by the excess production of ROS during oxidative phosphorylation. Note that although astrocytes mainly rely on glycolysis, there is evidence showing that astrocytes have oxidative phosphorylation, and it plays an important role in mitigating neuroinflammation and neurodegeneration^78^. We assume that the efflux rate is determined by the concentration gradient and the extent of mPTP opening. The calcium efflux rate is defined as shown in Eq. (8):

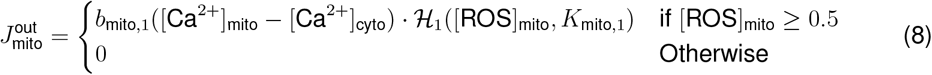

As for the influx via the mitochondrial membrane, we assume that it is only dependent on the cytosolic calcium concentration. Thus, we model it by Eq. (9):

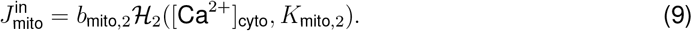

#### Calcium exchange across the plasma membrane

Calcium influx from the extracellular space into the cytoplasm occurs through both a small leak flux across the plasma membrane and ion channels. Previous studies, have modeled plasma membrane calcium channels in an IP_3_-dependent manner using a second-order Hill function based on IP_3_ concentration^10,79^. However, these models did not specify the ion channels involved, which limits their physiological relevance

In contrast, our model explicitly considers the involvement of voltage-gated calcium channels (VGCCs) and the *α*-amino-3-hydroxy-5-methyl-4-isoxazolepropionic acid (AMPA) receptors^80^. VGCCs are activated upon plasma membrane depolarization, meaning they open when the cytoplasmic concentration of positive ions increases, leading to a less negative membrane potential. This process facilitates calcium influx from the extracellular space. Additionally, AMPA receptors, a type of ionotropic glutamate receptor, open upon activation by extracellular glutamate, mediating calcium entry into the cell.

Our model of calcium flux via VGCCs is adapted from a model introduced in the literature^16^, which separately modeled four VGCC subtypes (T-type, L-type, N-type, and R-type) in astrocytes Given our focus on cortical astrocytes in mice, we relied on gene expression data^40,57^ and retained only the T-type and L-type channels, as they exhibit higher expression levels. In the model from the literature^16^, calcium flux through VGCCs primarily depends on intracellular and extracellular calcium concentrations, with the assumption that membrane potential and extracellular calcium levels remain constant. This assumption aligns with findings from the literature^3^, where patch-clamp recordings indicated that somatic membrane potential changes by less than 3 mV upon stimulation, despite more substantial, pathway-specific changes occurring in perisynaptic astrocyte processes. Since our model focuses on temporal dynamics, we have adopted the assumption of a constant membrane potential. Moreover, we incorporated the regulatory effect of PKA on VGCCs, as reported in the literature^81^, where PKA serves as an agonist for these channels.

For AMPA receptor modeling, we adopted the three-state Markov chain model from the literature^82^, which characterizes the AMPA receptor states as desensitized, closed, and open. In our model, [AMPA] ∈ [0, 1] represents the fraction of AMPA receptors that are activated at any given time, with this activation being primarily dependent on extracellular glutamate concentration.

Calcium influx from the extracellular space is described by Eq. (10), where the first term represents calcium flux via VGCCs, the second term captures flux via AMPA receptors, and the third term accounts for baseline flux across the membrane. In this equation, *v*_AMPA_ denotes the maximum calcium flux through AMPA receptors, *z* is the valence of calcium ions, *F* is the Faraday constant, *V*_ast_ represents the astrocyte volume, and *I*_VGCC_ is the electrical current through VGCCs. Details about the modeling of VGCC and AMPA can be found in the section titled “Receptor-mediated calcium entry and intracellular signaling”.

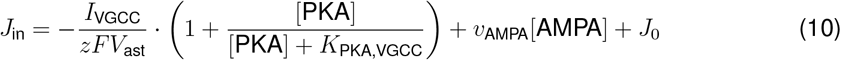

For calcium efflux to the extracellular space, a simple linear model with a coefficient of 0.5 was introduced to describe the flux from the cytoplasm to the extracellular space^83^. However, since our model features activated cPKC as a key factor contributing to the refractory period of astrocytic calcium, we additionally consider the excitatory effects of cPKC on the calcium ATPase transporter of the plasma membrane (PMCA) and the sodium calcium exchanger (NCX)^84^, which facilitate calcium extrusion from the cytoplasm to the extracellular space. Specifically, activation of cPKC after an increase in cytosolic calcium levels accelerates calcium efflux through PMCA and NCX. By integrating the model from the literature^83^ with our considerations of cPKC, we model the calcium flux from the cytoplasm to the extracellular space as described in Eq. (11):

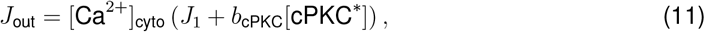

where we use [cPKC^∗^] ∈ [0, 1] to represent the percentage of cPKC being activated by calcium and DAG.

### Behavior-derived molecular inputs and signal construction

In this section, we outline our approach to derive the three input molecular signals to our virtual astrocyte model: norepinephrine (NE), dopamine (DA), and glutamate. Since direct measurements were not possible in our experiment, we inferred their levels based on the mouse’s run velocity and external stimuli, including visual cues and rewards.

#### Norepinephrine signal derivation

Run-evoked NE activity has been shown to strongly correlate with astrocytic calcium signals, as revealed by the GRAB_NE_ sensor^52^. Because NE promotes calcium release through the IP_3_-mediated pathway, this close temporal correlation suggests that astrocytic calcium dynamics may, in part, reflect filtered NE activity.

Previous studies have demonstrated that astrocytic calcium signals can be effectively modeled as leaky integrators of behavioral variables such as neuronal firing rate, paw movement, or pupil diameter^12^. Among these, paw movement showed the strongest correlation with astrocytic calcium activity. In our behavioral paradigm, running velocity serves as the analog of paw movement. Therefore, by extension, if calcium signals can be represented as a leaky integration of locomotor activity, and if NE and calcium are in phase, it is reasonable to model NE itself as a leaky integrator of run velocity.

Accordingly, we defined the NE signal *N* (*t*) as Eq. (12):

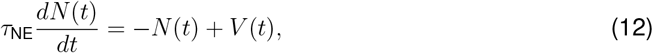

where *V* (*t*) is the spline-interpolated running velocity and *τ*_NE_ is the integration time constant governing the temporal smoothing of NE relative to velocity. The analytical solution to Eq. (12) is:

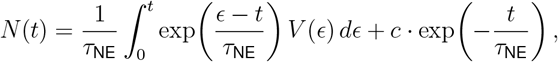

where *c* is the initial NE concentration. This formulation captures the slow, cumulative dynamics of NE release driven by locomotion, providing a biologically plausible intermediary between behavioral state and astrocytic calcium signaling.

#### Dopamine signal derivation

DA release in the cortex has been associated with both locomotion and reward processing. Electrical stimulation of dopaminergic projections from the ventral tegmental area to the motor cortex reinforces locomotor behavior and enhances reward-driven performance^54^. Motivated by these findings, our model represents DA inputs as two independent components: a run-evoked signal and a reward-evoked signal. This separation allows us to account for the distinct yet occasionally overlapping behavioral contexts in which cortical DA release occurs.

Experimental evidence shows that DA concentration increases sharply following run/reward onset and decays slowly, consistent with the kinetics observed using the fluorescent dopamine indicator dLight1.2 in the motor cortex^55,56^. To capture this biphasic profile, each DA transient is modeled as the product of two Hill functions: one governing the rising phase and the other (reversed) governing the decay. Suppose that a reward delivery or the onset of running occurs at time *t*_*i*_. Then the DA response to this event, denoted *D*_*i*_(*t*), can be expressed as Eq. (13):

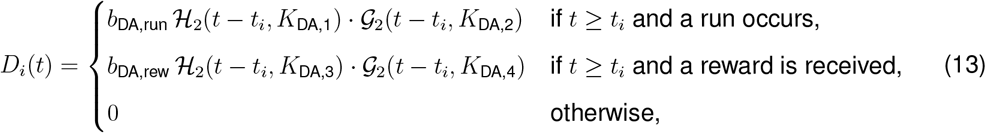

where

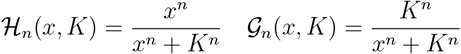

are the Hill function and its reversed form. The parameters *b*_DA,run_ and *b*_DA,rew_ are scaling coefficients controlling the amplitude of run-evoked and reward-evoked DA release, respectively, while *K*_DA,1_–*K*_DA,4_ define the steepness and duration of the rise and decay phases. The total DA concentration over time is shown in Eq. (14), which is a linear superposition of all individual DA responses, assuming *n*_*D*_ run or reward events:

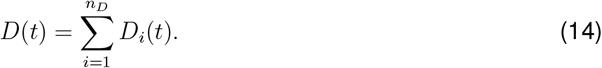

This formulation captures the temporally asymmetric rise-and-fall dynamics of DA release and provides a flexible framework for simulating cortical DA signals arising from both motor and reward-related events.

#### Glutamate signal derivation

Glutamate signaling differs fundamentally from NE and DA in that glutamate not only acts on astrocytic membrane receptors but is also taken up into astrocytes via excitatory amino acid transporters (EAATs), contributing directly to cellular metabolism. Moreover, given the high density of glutamatergic neurons in the motor cortex, we explicitly modeled the glutamate dynamics between neurons and astrocytes. Once transported into astrocytes, glutamate is converted into glutamine, which is subsequently shuttled back to neurons, completing the glutamate–glutamine metabolic cycle.

Consistent with experimental observations that extracellular glutamate levels increase during locomotion and are further modulated by sensory inputs^53,58^, we modeled neuronal glutamate release as impulses triggered by either running or visual stimulation. Each glutamate impulse exhibits a rapid rise and slower decay, and thus it is modeled in the similar way as DA, as shown in Eq. (15):

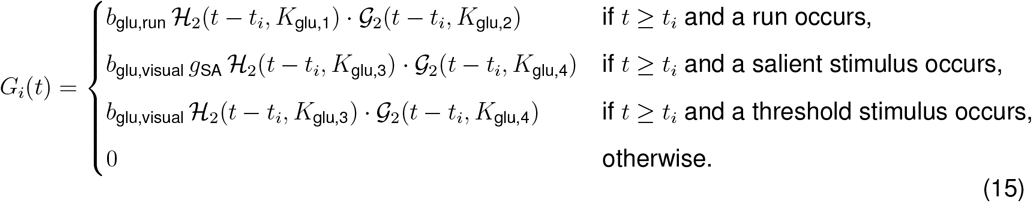

Here, *b*_glu,run_ and *b*_glu,visual_ denote the amplitude coefficients for locomotion- and stimulus-evoked glutamate release, respectively; *g*_SA_ represents the gain factor for salient stimuli; and *K*_glu,1_–*K*_glu,4_ specify the rise and decay kinetics. The total neuronal glutamate concentration is obtained by summing over all *n*_*G*_ events, as shown in Eq. (16):

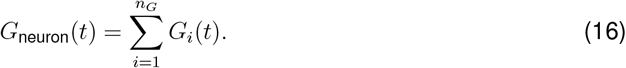

To capture neurotransmitter trafficking between compartments, we modeled the temporal dynamics of glutamate in the synaptic cleft (*G*_cleft_), in the astrocyte cytoplasm (*G*_ast_), and the resulting astrocytic glutamine pool (*G*_gln_) as shown in Eqs. (17), (18), (19):

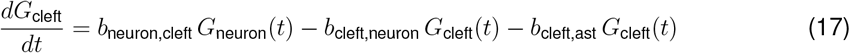

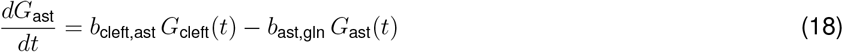

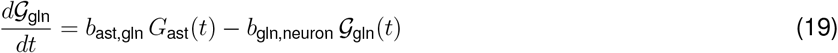

This formulation captures the continuous exchange of glutamate and glutamine between neurons and astrocytes, allowing the model to reproduce both receptor-mediated calcium signaling and transporter-driven metabolic coupling. The use of compartmental glutamate representation also enables region-specific extensions, such as inclusion of calcium-permeable AMPA receptor contributions or developmental re-expression of mGluR5.

### Receptor-mediated calcium entry and intracellular signaling

In this section, we model five membrane receptors or channels: three GPCRs (Gq, Gs, Gi), AMPA receptors, and VGCCs. Throughout this section, we frequently use Hill functions to represent saturating biochemical relationships. These functions generalize the classical Michaelis–Menten form when the Hill coefficient *n* = 1.

#### G-protein coupled receptor modeling

Activation of Gq-GPCR triggers phospholipase C (PLC), which hydrolyzes phosphatidylinositol 4,5-bisphosphate (PIP_2_) into inositol trisphosphate (IP_3_) and DAG, initiating calcium signaling through the IP_3_-mediated pathway^35^. In contrast, activation of Gs-GPCR stimulates adenylyl cyclase (AC), which converts adenosine triphosphate (ATP) into cyclic adenosine monophosphate (cAMP). Elevated cAMP levels then activate PKA, which not only facilitates calcium release from the ER via the IP_3_ receptor (IP_3_R) channel but also phosphorylates transcription factors involved in regulating gene expression^85^. In contrast to Gs-GPCR, Gi-GPCR inhibits the activities of AC, which then inhibits the cAMP-PKA pathway Meanwhile, AMPA receptors, which are ionotropic glutamate receptors, open in response to glutamate released from neurons, thereby mediating calcium influx from the extracellular space into the cytoplasm. This increase in cytosolic calcium can further trigger calcium-induced calcium release (CICR) when IP_3_ is present.

For each receptor, we assume that the total number of receptors is conserved. Let [GqGPCR^∗^], [GsGPCR^∗^], [GiGPCR^∗^], and [AMPA^∗^] represent the fractions of each receptor type that are activated. Then the time derivatives of these activated receptors follow a generic form, described by Eq. (20):

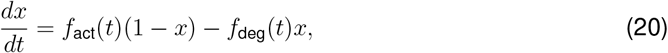

where *x* represents the activated percentage of receptors, *f*_act_(*t*) represents the time-variant activation factor that depends on agonists, and *f*_deg_(*t*) represents the time-variant degradation factor that depends on antagonists.

In mouse astrocytes, genes for *α*-adrenergic receptors and metabotropic glutamate receptor 5 (mGluR5) are found, and these receptors function as Gq-GPCRs^40,57^. Additionally, heteromers formed by D1-like and D2-like dopamine receptors are also classified as Gq-GPCRs^86^. The activation of Gq-GPCRs is modeled as a linear combination of extracellular norepinephrine (NE, represented by *N* (*t*)), dopamine (DA, represented by *D*(*t*)), and glutamate (represented by *G*_cleft_(*t*)), all of which are time-variant. The degradation of Gq-GPCRs is assumed to depend on the activated conventional protein kinase C (cPKC) as well as other factors. Eq. (21) describes the dynamics of Gq-GPCR activation and degradation:

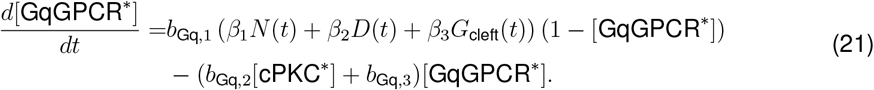

Mouse astrocytes also express genes for *β*-adrenergic receptors, as well as D1 and D5 dopamine receptors, which are classified as Gs-GPCRs^40,57^. Therefore, we model the activation of Gs-GPCRs as a linear combination of norepinephrine (NE) and dopamine (DA). The degradation of Gs-GPCRs is assumed to be mediated by PKA and other factors, with degradation rates being linearly dependent on the number of active receptors. Eq. (22) represents the dynamics of Gs-GPCRs:

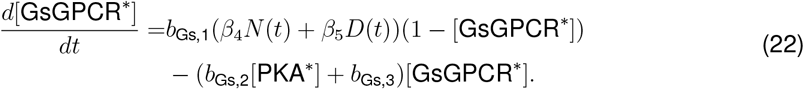

Besides Gq and Gs-GPCRs, mouse astrocytes also have Gi-GPCRs such as D2-like DA receptors, metabotropic glutamate receptor 3 (mGluR3), and *α*2-adrenergic receptors^40,57^. Thus, in our model, all three input agonists activate the Gi-GPCRs, and their degradation depends on PKA and cAMP. Importantly, recent experimental evidence demonstrated that NE, acting through *α*2-adrenergic receptors (Gi-GPCRs), can gate astrocytic responses to dopamine by lowering cAMP levels, thereby alleviating the inhibitory effect of cAMP/PKA on D2-like receptors and enhancing D1–D2 heteromer formation, which in turn activates Gq-coupled calcium signaling pathways^4^. To capture this effect, our model explicitly implements NE-dependent inhibition of the cAMP–PKA pathway as a mechanism for gating dopaminergic signaling. Eq. (23) represents the dynamics of Gi-GPCRs:

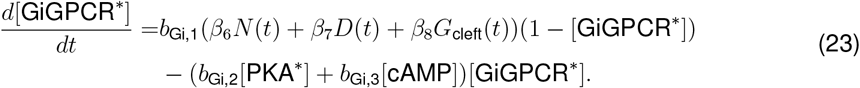

#### AMPA receptor modeling

Mouse astrocytes also highly express genes for AMPA receptors, a class of ionotropic glutamate receptors that are activated by extracellular glutamate and mediate calcium influx from the extracellular space. For modeling AMPA receptor activity, we employ the Desensitized-Closed-Open (DCO) model introduced in the literature^82^, which uses a three-state Markov chain to describe AMPA receptor dynamics. Let **p**(*t*) = [*p*_*D*_, *p*_*C*_, *p*_*O*_] be the probability distribution of the state of an AMPA receptor at time *t*. Then the dynamics of **p**(*t*) are described by:

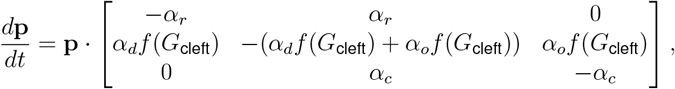

where

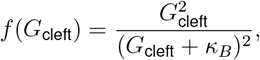

and *G*_cleft_ represents the extracellular glutamate level. Here, *α*_*o*_ is the opening rate, *α*_*c*_ is the closing rate, *α*_*d*_ is the desensitization rate, *α*_*r*_ is the re-sensitization rate, and *κ*_*B*_ is the dissociation constant. Based on this continuous Markov chain shown in Eq. (24), the dynamics of [AMPA] are given by:

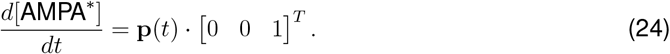

#### HVGCC modeling

The VGCCs play a crucial role in mediating calcium influx from the extracellular space into the cytoplasm. When modeling the calcium current via VGCC (*I*_VGCC_ in Eq. (10)), the widely accepted generic form is given by Eq. (25)^16,48,87,88^:

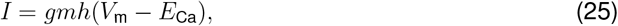

where *g* is the membrane conductance (in pS), *m* is the channel activation factor, *h* is the channel inactivation factor, *V*_m_ is the membrane potential (in mV), and *E*_Ca_ is the Nernst potential of calcium (in mV). VGCC dynamics are typically modeled under two assumptions: either the membrane potential (*V*_m_) is constant while the Nernst potential (*E*_Ca_) varies, as in astrocyte models^16^, or *V*_m_ is variable while *E*_Ca_ is constant, as in models of pancreatic *β* cells^48,87,88^. According to the literature^3^, the membrane potential in astrocyte somata changes by less than 3 mV upon stimulation, whereas peripheral astrocyte processes show significant changes. In our case, since the model follows a mean-field approach and describes calcium dynamics across the entire cell, we assume *V*_m_ is constant while *E*_Ca_ is variable.

Among the four VGCC subtypes (T-type, L-type, N-type, and R-type), only T-type and L-type channels are strongly expressed in mouse astrocytes. Therefore, we retain only the T-type and L-type components from the model in the literature^16^. The activation factor *m* and the inactivation factor *h* are modeled as Eqs. (26), (27):

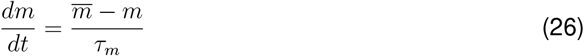

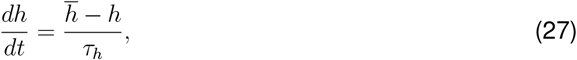

where 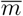 and 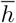 represent the steady-state values that depend on the membrane potential, and *τ*_*m*_ and *τ*_*h*_ are time constants that control the rates of convergence for *m* and *h*, respectively, both of which are also membrane potential-dependent.

The calcium current via the T-type VGCC is modeled by Eq. (28):

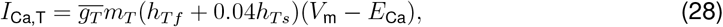

where *m*_*T*_, *h*_*Tf*_, and *h*_*Ts*_ follow the generic dynamics captured in Eqs. (26), (27). The Nernst potential, *E*_Ca_, is modeled by Eq. 29:

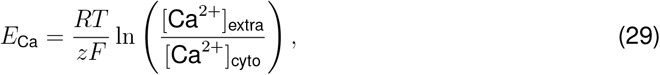

where *R* is the ideal gas constant, *T* is temperature, *z* is the valence of calcium ion, *F* is the Faraday constant, and [Ca^2+^]_extra_ is the extracellular calcium concentration (constant).

Similarly, the calcium current via the L-type VGCC is modeled by Eq. (30):

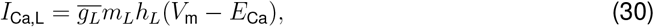

where *m*_*L*_ follows the generic dynamics in Eq. (26), while *h*_*L*_ does not follow Eq. (27), but is dependent on the cytosolic calcium, as shown in Eq. (31):

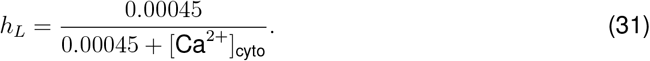

### Modeling intracellular molecular signaling pathways

In this section, we introduce the modeling of intracellular molecular signals other than calcium. We categorize these signals into three groups: IP_3_ pathway-related, mitochondria-related, and cAMP-PKA-related. These categories correspond to the three circuits in our model. Similar to the previous section, we frequently use Hill functions to represent saturating biochemical relationships.

#### IP_3_ pathway-related signals

We assume that the total amount of phospholipase C (PLC) is conserved. PLC is activated by active Gq-GPCR and deactivated by active conventional protein kinase C (cPKC) and possibly other factors^35^. The model is presented in Eq. (32), which follows the generic format of Eq. (20):

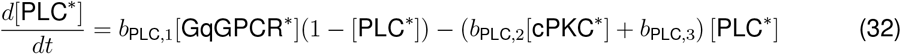

When PLC is activated, it hydrolyzes PIP_2_, producing both IP_3_ and DAG at the same rate. For simplicity, we do not model the dynamics of PIP_2_ directly but instead quantify the effect of PLC using a first-order Hill function, as shown in the initial part of Eqs. (33), (34). For the degradation of IP_3_ and DAG, we refer to the literature^35^:

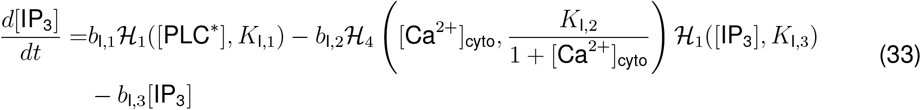

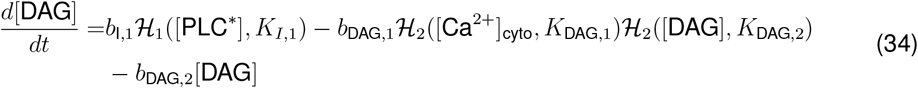

When cytosolic calcium levels rise, cPKC is activated by binding to one calcium ion and one DAG molecule. Upon activation, cPKC translocates to the plasma membrane, where it inhibits PLC and Gq-GPCR, forming a negative feedback loop. We assume that the total amount of cPKC is conserved. Given that activation involves two ligand-binding processes, we model it using two first-order Hill functions. For degradation, due to the limited references available, we represent it with a linear term. The dynamics of cPKC are shown in Eq. (35), which aligns with the generic form in Eq. (20):

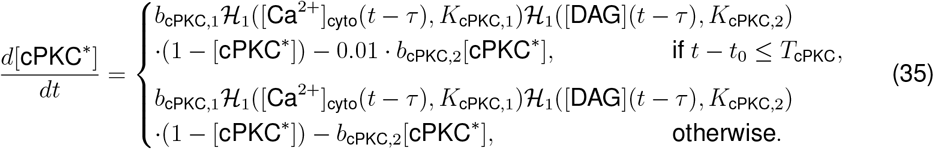

where *T*_cPKC_ is the cPKC preserving time, and *t*_0_ is the most recent time that calcium reaches a certain threshold (defined by [Ca^2+^]_threshold_):

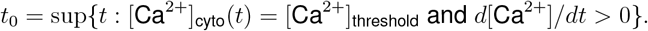

#### Mitochondria-related signals

Since we assume that NE is triggered by the mouse’s locomotion, glutamate is triggered by both locomotion and visual stimuli, and dopamine is triggered by locomotion and reward, we model the level of ATP, which reflects the level of metabolism, as a linear combination of NE and glutamate. This relationship is shown in Eq. (36):

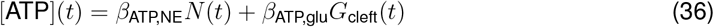

Regarding ROS, since it is a product of oxidative phosphorylation on the inner mitochondrial membrane, we assume its production depends on glutamate with a time delay, *τ*. When ROS accumulates within the mitochondria and surpasses a certain threshold, the mPTP opens, allowing ROS to diffuse into the cytoplasm. In the cytoplasm, ROS sensitizes IP_3_R2 and is degraded by superoxide dismutases. Thus, we model ROS dynamics in both mitochondria and cytoplasm, as shown in the Eqs. (37) (38):

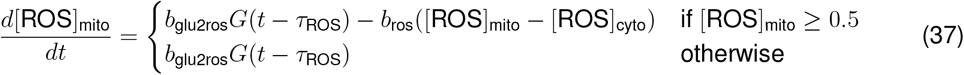

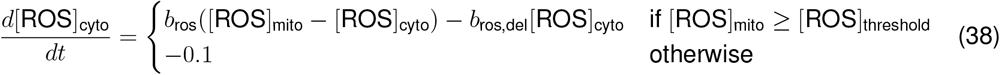

#### cAMP-PKA-related signals

In the cAMP-PKA pathway, not only does PKA independently trigger calcium release from the ER^48^, but PKA itself is also an important signaling molecule that causes many downstream effects. In this subsection, we present a simplified model of the calcium-cAMP-PKA circuit inspired by the literature^48^.

We assume that the total amount of AC is conserved, and it is activated by active Gs-GPCR and deactivated by PKA. The modeling is shown in Eq. (39):

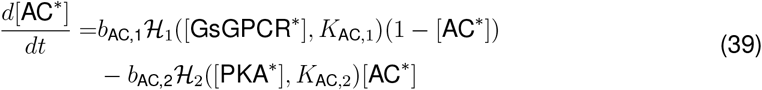

For cAMP, which is produced from ATP via the catalysis of AC, we assume that production is linearly dependent on the active AC, and degradation is dependent on active phosphodiesterase (PDE) and other factors. The modeling is shown in Eq. (40):

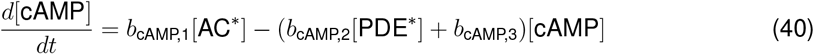

For PKA, we also assume its total amount is conserved. It is activated by cAMP and degraded by some unknown factors that are linearly dependent on the amount of active PKA. The modeling is shown in Eq. (41):

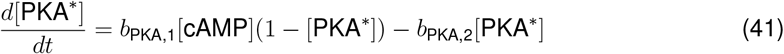

For PDE, we also assume its total amount is conserved. It is activated by cytosolic calcium and degraded by some unknown factors that are linearly dependent on the amount of active PDE The modeling is shown in Eq. (42):

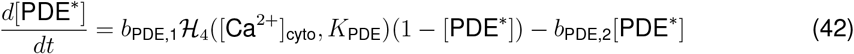

## Appendix S2

### Joint regulation of calcium dynamics by IP_3_, ATP, and cytosolic calcium

In this section, we model the marginal effects of cytosolic calcium, IP_3_, and ATP on the open probability of IP_3_R2, as a supplementary elaboration on Eq. (5). The unknown parameters are estimated using experimental data from the literature^45^, which provides open probability values at various concentrations of the three chemicals. The functional forms of the marginal effects are designed based on the curve shapes shown in the literature^45^.

Motivated by the characteristic shapes of the IP_3_R2 curves reported in the literature^45^, we modeled the marginal effect of [Ca^2+^]_cyto_ using a log-Gaussian function, as shown in Eq. (43), where the mean and standard deviation are treated as unknown parameters:

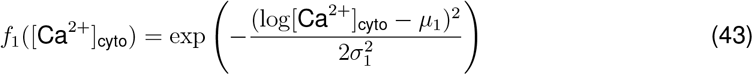

Since ATP exhibits an excitatory effect on IP_3_R2 at low concentrations and an inhibitory effect at high concentrations^46^, we also use a log-Gaussian function to model its marginal effect. Curves for IP_3_R2 in the literature^45^ show that when the IP_3_ concentration is high (10 *µ*M), the effects of both high ATP (5 mM) and low ATP (0.01 mM) are similar. However, at low IP_3_ concentration (1 *µ*M), high ATP leads to a significantly higher open probability compared to low ATP. Thus, we model the marginal effect of ATP using Eq. (44):

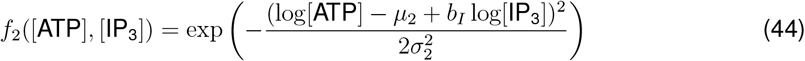

For the effect of IP_3_, a common practice is to use a Hill function. Curves for IP_3_R2 in the literature^45^ show that when ATP is high (5 mM), there is little difference between high and low IP_3_ levels. However, when ATP is low (0.01 mM), high IP_3_ results in a significantly higher open probability. Thus, we model the marginal effect of IP_3_, dependent on ATP, with Eq. (45):

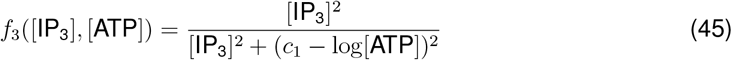

Define ***θ*** = [*µ*_1_, *σ*_1_, *µ*_2_, *σ*_2_, *b*_*I*_, *c*_1_]^*T*^ as the vector of unknown parameters in the marginal effect functions. Then the joint effect of cytosolic calcium, ATP, and IP_3_ on IP_3_R2 open probability can be expressed as Eq. (46):

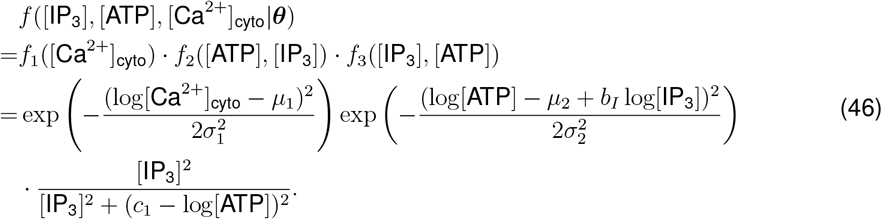

Using data points from the literature^45^, we define the loss function as the root mean squared error (RMSE):

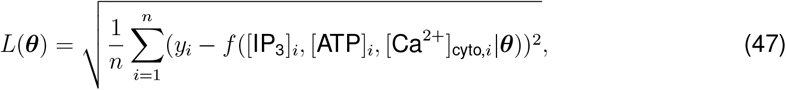

where *y*_*i*_ is the real open probability. The first-order gradient of the loss in Eq. (47) is:

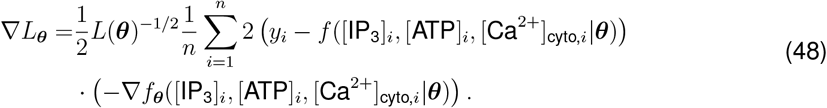

The gradient of *f*, i.e., ∇*f*_***θ***_, can be easily obtained in the following way:

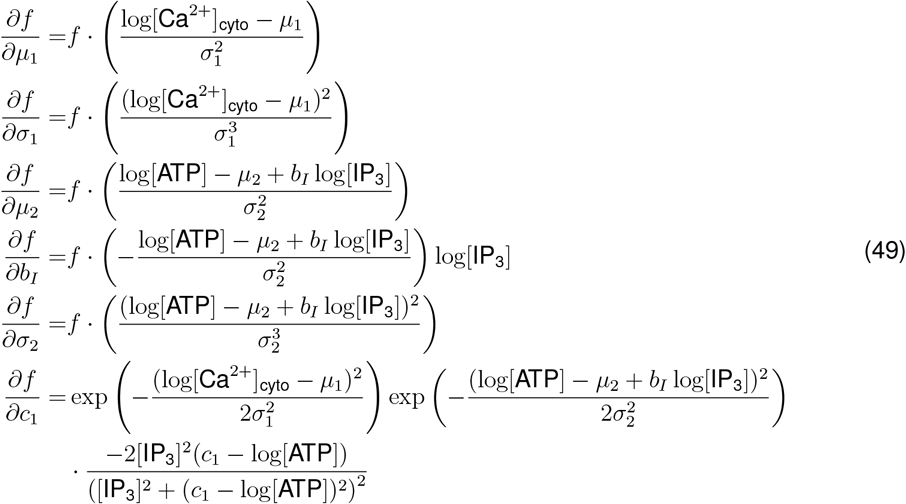

Using the first-order gradient, we can approximate the optimal value ***θ***^∗^ by the stochastic gradient descent (SGD) method. We provide the loss with respect to time and fitted open probabilities in Figure S2I-J.

## Appendix S3

### Derivation of calcium upper bound from fluorescence signals

In this section, we describe the procedure used to compute the minimum fluorescence, *F*_min_, which is the fluorescent intensity in the absence of calcium ions. Once *F*_min_ is determined, we combine it with the known parameter ranges of GCaMP6f and the observed fluorescence baseline (i.e., *F*_0_) and Δ*F/F*_0_ values to derive an upper bound for the cytosolic calcium concentration in our data. For the dynamic range of the GCaMP6f indicator, i.e., *F*_max_*/F*_min_, since the values provided in literature were measured in vitro^69^, which may not necessarily represent the *in vivo* case, we do not adopt the values of *F*_max_*/F*_min_ shown in the literature, but instead assume that the maximum fluorescent intensity occurs at the maximum value of Δ*F/F*_0_ among all calcium events detected by AQuA2 in our datasets. Assume there are *N* detected calcium events in all calcium recordings, and let

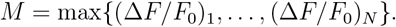

Then the relationship between calcium concentration and fluorescence intensity is modeled by the following Hill equation^70^:

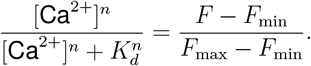

In this equation, *F* denotes the fluorescence intensity corresponding to a specific calcium concentration, *F*_min_ represents the fluorescence in the absence of calcium, and *F*_max_ corresponds to the fluorescence at saturating calcium levels. Meanwhile, the parameters *n, K*_*d*_ for GCaMP6f are known to lie within the ranges *n* ∈ [2.17, 2.37], and *K*_*d*_ ∈ [0.361, 0.389]^69^. However, in our imaging data, the cytosolic calcium in astrocytes is never truly zero but instead remains at approximately 50–150 nM under resting conditions^34^. Therefore, the measured baseline intensity, denoted by *F*_0_, reflects the fluorescence corresponding to these low calcium levels rather than a true zero-calcium state. Thus, we used the following equation:

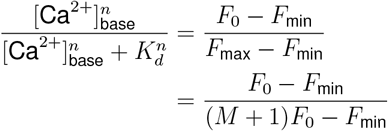

which yields

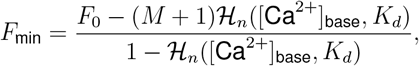

where [Ca^2+^]_base_ ∈ [0.05, 0.15] (in *µ*M) is the calcium concentration under resting conditions and ℋ_*n*_([Ca ]_base_, *K*_*d*_) is the n-th order Hill function. Then, the peak value of calcium corresponding to a calcium event with baseline fluorescence being *F*_0_ and proportional change of fluorescence being Δ*F/F*_0_ satisfies

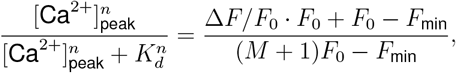

and thus

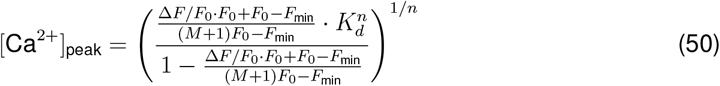

We set *M* = 5.8 because the maximum observed Δ*F/F*_0_ is 5.8 (Figure S4B). To determine the overall upper bound for cytosolic calcium across all detected events, we performed a grid search over the parameter space. Specifically, we varied *K*_*d*_ from 0.361 to 0.389 in increments of 0.001, *n* from 2.17 to 2.37 in increments of 0.01, and [Ca^2+^]_base_ from 0.05 to 0.15 *µ*M in increments of In essence, the grid search aimed to find

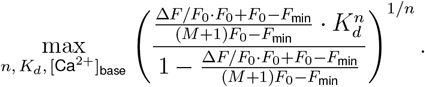

Since we assumed the maximum fluorescence *F*_max_ occurs when Δ*F/F* reaches a maximum, the derived [Ca^2+^]_peak_ for the calcium event having the maximum Δ*F/F* is infinity. We provide the histogram of the grid-search results, after excluding the infinity values, in Figure S4B. We observe that most data points in the histogram are less than 0.5, indicating that, in most cases, calcium peak values during calcium activity are less than 0.5 *µ*M. The upper bound for the intracellular calcium signal, after adjusting for *F*_min_, is determined to be 2.618 *µ*M, corresponding to the parameter set *K*_*d*_ = 0.389, *n* = 2.17, and [Ca^2+^]_base_ = 0.15. However, when [Ca^2+^]_base_ = 0.15*µ*M, since the volume of the ER is 15%-70% of the cytoplasm^35^ and the calcium in the ER is at least 1000 times that in cytoplasm^33^, the cytosolic calcium should be at least 22.5 *µ*M to deplete the ER, which is far beyond 2.618 *µ*M that we obtained. Therefore, we conclude that the ER is unlikely to be depleted in our experiments.

## Appendix S4

### Model parameter optimization and fitting procedures

In our calcium signaling model, each equation includes parameters that must be specified. While some of these parameters can be adopted directly from the literature, others require optimization tailored to our experimental conditions. We denote the set of parameters to be optimized as **Θ**, and refer to a particular choice of parameter values as a configuration. The quality of any given configuration is assessed by measuring the similarity between experimentally recorded calcium signals and those simulated by the model using that configuration. To systematically determine the optimal configuration, we first define metrics to quantify signal similarity and then apply an appropriate optimization algorithm.

As described in the **Results**, model performance is assessed using five complementary metrics: root mean squared error (RMSE), activity-dependent RMSE, Pearson correlation coefficient, cosine similarity, and normalized dynamic time warping (DTW) distance. These metrics quantify different aspects of similarity between the real and predicted calcium signals:

- **RMSE** measures the average pointwise deviation between the predicted and real signals across the entire trace. Because all traces are normalized to the range [0.05, 1], RMSE lies in range [0, 0.95], with values closer to 0 indicating higher accuracy.
- **Activity-dependent RMSE** measures the same deviation but restricted to time points in which the recorded signal exhibits calcium activity (i.e., values above a threshold), thereby capturing performance during physiologically meaningful transients. As with RMSE, its range is [0, 0.95] under signal normalization.
- **Correlation** quantifies temporal synchrony by measuring how well deviations from the mean align across traces. Its range is [−1, 1], though values in this study typically fall within [0, 1], with values near 1 indicating high synchrony.
- **Cosine similarity** treats each trace as a vector in R^*T*^ and measures the cosine of the angle between them. For nonnegative normalized signals, the cosine similarity lies strictly in [0, 1], with values near 1 indicating strong agreement.
- **Normalized DTW distance** measures temporal alignment by computing the dynamic time warping cost and dividing by the trace length, producing a scale-free measure. Values in [0, 0.05] correspond to an almost identical match, 0.05–0.10 indicate a good match, 0.10–0.20 reflect a moderate mismatch, and values exceeding 0.20 indicate substantial temporal or shape differences.

To cast parameter fitting as a minimization problem, we negate the correlation and cosine similarity scores so that lower values across all metrics indicate better performance. Suppose we evaluate **Θ** on *n* calcium recordings, and let **Y**_*i*_ denote the real calcium trace from the *i*-th recording and 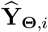 the corresponding simulated trace, then the five objective functions are:

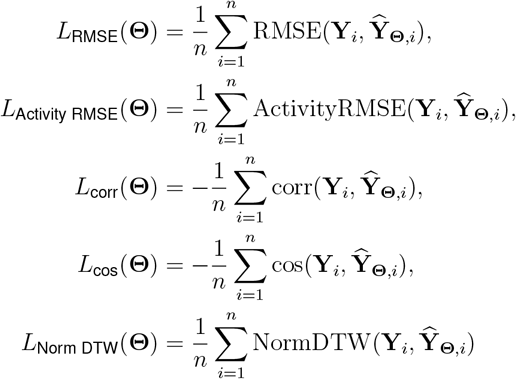

For two time series **X, Y** ∈ ℝ^*T*^ with *k* calcium activities occurring over disjoint time intervals 𝒜_1_, …, 𝒜_*k*_ ⊆ *{*1, …, *T}*, let 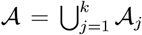 denote the full set of activity time points. Then the metrics are defined as follows:

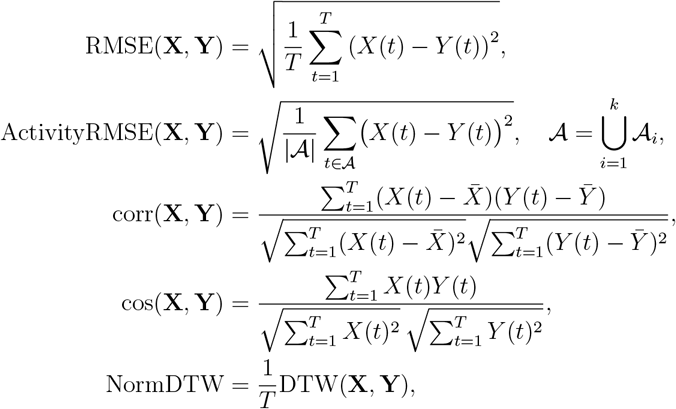

where 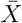 and 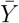 are sample means, |𝒜 | is the number of active time points, and DTW(·,·) denotes the standard dynamic time warping cost, computed using MATLAB’s built-in dtw function. In practice, the disjoint activity intervals 𝒜_1_, …, 𝒜_*k*_ used in the activity-dependent RMSE are determined from the real calcium trace by thresholding at baseline+10% of the calcium peak amplitude. Each objective function therefore reflects a distinct aspect of trace similarity, and the multi-objective optimization seeks parameter configurations that perform well across all five criteria.

In single-objective optimization problems, the objective is a scalar quantity, making comparisons via greater-than (*>*) and less-than (*<*) straightforward. However, in the multi-objective setting, we replace these scalar comparisons with the concept of domination. A commonly used definition is Pareto-domination, as discussed in the literature^89^ (Chapter 28). For two configurations **Θ**_1_ and **Θ**_2_, let *L*_*i*_ denote the *i*-th objective function for *i* = 1, …, *M*. Then, **Θ**_1_ is said to Pareto-dominate **Θ**_2_ if the following two conditions are satisfied:

1. ∀*i* ∈ *{*1, 2, …, *M}, L*_*i*_(**Θ**_1_) ≤ *L*_*i*_(**Θ**_2_)

2. ∃*j* ∈ *{*1, 2, …, *M}* such that *L*_*j*_(**Θ**_1_) *< L*_*j*_(**Θ**_2_)

In other words, configuration **Θ**_1_ is no worse than **Θ**_2_ in all objectives and strictly better in at least one.

Given the five evaluation metrics and the Pareto-domination condition, we selected the genetic algorithm (GA) as the optimization method, as it has been empirically found to be effective in optimizing single-neuron models^71,90^. Additional discussion can be found in the literature^89^ (Chapter 28). The GA is a population-based stochastic optimization technique inspired by the process of natural selection. It begins with an initial population of randomly generated parameter configurations, which are like chromosomes in biology and evolve over successive generations. In each generation, candidate configurations are evaluated using a fitness function, which in our case is a multi-objective loss based on similarity between real and predicted calcium signals. The best-performing configurations are selected and undergo genetic operations such as crossover (the recombination of parameter values in pairs of configurations) and mutation (random perturbation) to create a new population. Over time, this evolutionary process guides the population toward parameter sets that yield improved model performance. Below is the pseudocode for the GA on multi-objective optimization.

#### Algorithm 1

Multi-objective Genetic Algorithm for Parameter Optimization

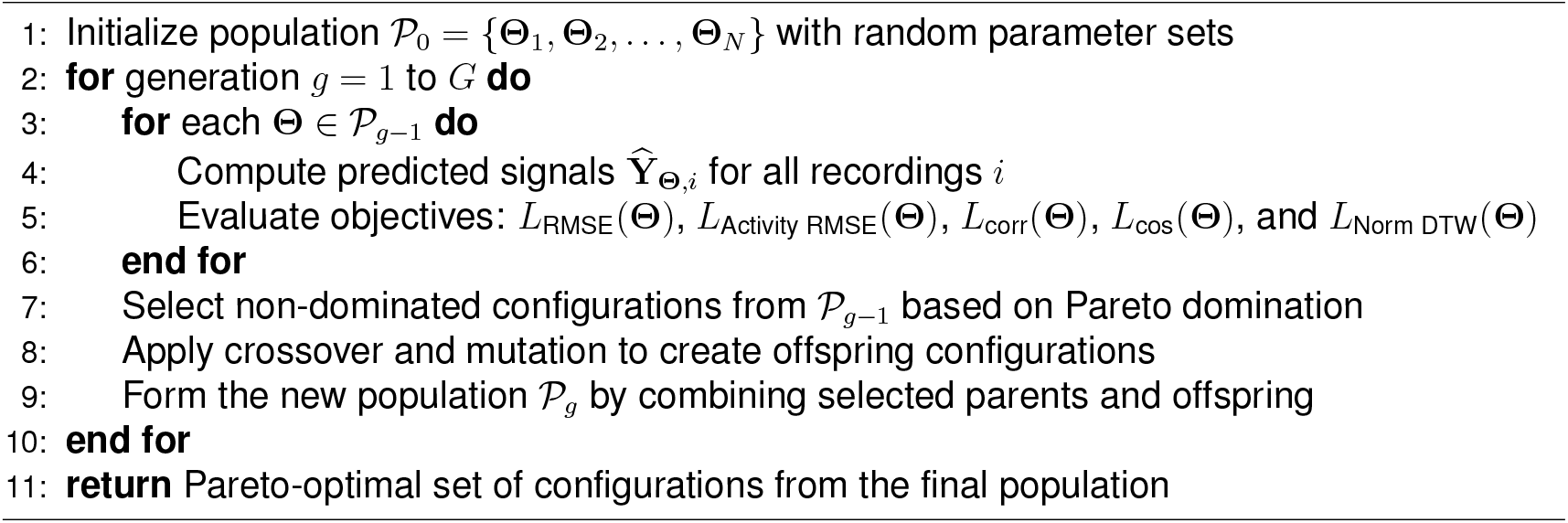

To ensure that the parameter optimization process generalizes well to new data and does not simply overfit to a specific subset of recordings, we employed a stratified five-fold cross-validation (CV) strategy in conjunction with the GA. Cross-validation is a common approach in machine learning used to test a model’s ability to make accurate predictions on data it has not seen before In a five-fold CV, the whole dataset—in our case, 35 calcium recordings—is split into five equally sized parts, or “folds”, each containing 7 recordings. To preserve consistency, the splitting was stratified so that each fold contained a similar proportion of recordings from each mouse. This step is crucial for avoiding biases introduced by mouse-specific patterns and ensures that each fold is representative of the full dataset.

In each round of the five-fold CV, one fold was set aside as a test set, while the remaining four folds (28 recordings) were used as a training set to perform GA optimization on the model parameters. Within the training folds, the GA was run to optimize the model parameters. The GA maintains a population of candidate parameter configurations and iteratively evolves them through biologically inspired operations—such as selection, crossover, and mutation—to search for configurations that improve model performance. Optimization was guided by the evaluation metrics described above.

After running the GA for a predefined number of generations on the training data, we selected one final parameter configuration from the last generation’s Pareto front—a set of solutions that represent trade-offs among the five objectives. The specific configuration chosen was the one with the lowest RMSE on the training data. This configuration was then applied to simulate calcium signals on the test fold, providing an independent evaluation of model performance.

This entire process was repeated five times, rotating the test fold each time, so that every recording was used exactly once for testing. As a result, we obtained five sets of training and test results. The model’s performance across all five test folds, measured using the same five metrics, closely matched its training performance (Figure S5). This consistency indicates that the optimization approach did not overfit to the training data and that the model generalized well to unseen recordings. By combining stratified cross-validation with evolutionary parameter optimization, we achieved a robust and unbiased assessment of the model’s generalizability.

## Appendix S5

### Linear mixed-effect model analysis

Our analysis of the real calcium signal data and run velocity data revealed that the inter-run interval (IRI) plays a central role in shaping the amplitude of the second calcium response (A_2_) (Fig. 4A, B). Quantifying the relationship between A_2_ and IRI allows us to estimate the timescale of cPKC degradation, in particular, the half-time decay of its refractory effect. Because the amplitude of the first calcium response (A_1_) may also modulate the refractory state of the cell and thereby confound the IRI–A_2_ relationship, we also included A_1_ and its interaction with IRI as covariates. To account for these factors, we employed a linear mixed-effects model (LME) with IRI, A_1_, and their interaction as fixed effects. Given that the dataset contained repeated trials from multiple animals, random intercepts were included for each mouse to capture between-mouse variability. Conceptually, the LME comprises two hierarchical levels: a within-mouse level that describes trial-to-trial effects, and a between-mice level that accounts for differences in baseline calcium responses across animals.

### Level 1 (within-mouse)

We model the amplitude of the second calcium response for the *i*-th trial from the *j*-th mouse as

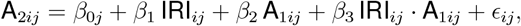

where A_1*ij*_ and A_2*ij*_ denote the amplitudes of the first and second calcium responses, respectively, IRI_*ij*_ is the inter-run interval, *β*_0*j*_ is a mouse-specific intercept (random effect), *β*_1_, *β*_2_, *β*_3_ are fixed-effect coefficients, and *ϵ*_*ij*_ ∼ 𝒩 (0, *σ*^2^) represents the within-mouse residual error.

### Level 2 (between-mice)

The mouse-specific intercept *β*_0*j*_ is modeled as

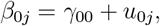

where *γ*_00_ is the overall average intercept (fixed effect) and *u*_0*j*_ ∼ 𝒩 (0, *τ* ^2^) is the mouse-specific deviation (random intercept).

### Combined model

Substituting the Level 2 model into Level 1 yields

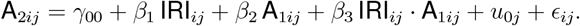

Before model fitting, the predictors were standardized (z-scored) so that coefficients could be directly compared across variables. The fitted LME revealed that the inter-run interval (IRI) was a highly significant predictor of A_2_. Specifically, the effect of IRI was positive (*β*_1_ = 0.16 ± 0.021, *t* = 7.54, p-value= 1.4 *×* 10^−12^), indicating that longer intervals led to larger second responses Interpreted on the standardized scale, this corresponds to an increase of 0.16 in A_2_ per one standard deviation of IRI. Converting back to raw units, where the standard deviation of IRI was 14.91 seconds, the slope is approximately 0.010 A_2_ units per second of IRI (Figure S3A).

By contrast, A_1_ and the IRI *×* A_1_ interaction were not significant (p-value = 0.28 and 0.34, respectively). This indicates that once IRI is accounted for, A_1_ does not explain additional variation in A_2_, nor does it modify the relationship between IRI and A_2_. Therefore, these terms were not retained for interpretation in the final model. Meanwhile, the insignificance of A_1_ and the IRI *×* A_1_ interaction further supports our conclusion that the ER is unlikely to deplete during calcium activities (as detailed in Appendix S3). If ER depletion occurred following the first calcium response, then A_1_ would be expected to significantly influence the amplitude of A_2_, which was not observed.

Random effects analysis supported the inclusion of random intercepts to account for between-mice differences in baseline A_2_, but did not support random slopes for IRI. A likelihood ratio test comparing the random-intercept model with the random-intercept–and–slope model yielded *χ*^2^(2) = 0.96, p-value= 0.62, indicating that allowing each mouse to have its own IRI effect did not improve model fit compared to the simpler random-intercept model. Thus, the positive IRI–A_2_ relationship is consistent across animals, with modest variability only in baseline response amplitude. The fitted regression lines for the population effect and for individual mice are shown in Figure S3A, B. According to the fitted line, the amplitude of the second calcium response starts at about 0.34 when the inter-run interval is 0 second. The refractory effect is fully degraded by roughly 63 seconds, when the second-event amplitude reaches 1. By linear interpolation, the response reaches half of this recovery at around 32 seconds, which can be interpreted as the half-time of cPKC degradation. The estimated times of full and half degradation of the refractory effect are consistent with values reported in the literature^8,9^, strengthening our confidence in the LME model.

These timescales can be further interpreted as the sum of sequential processes. For the case of full degradation at approximately 63 seconds, the duration can be attributed to: about 10 seconds from run onset to the calcium peak, 15 seconds for cPKC to reach its peak following the calcium rise, 20 seconds during which cPKC remains above the inhibitory threshold and effectively suppresses calcium responses, and 15 seconds for cPKC to degrade back to baseline For the case of half degradation at approximately 32 seconds, the timeline likely reflects 10 seconds for the calcium response to peak, 15 seconds for cPKC activation to peak, and about 7 seconds of partial cPKC degradation to reduce its inhibitory effect on calcium by half. This 7-second estimate matches the value reported in the main text, ensuring consistency between the detailed breakdown here and the summary provided in the **Results**. Together, these breakdowns provide a mechanistic interpretation of the LME-derived full and half-time degradation estimates

## Appendix S6

**Table 1.** Modeled signaling molecules, receptors, and calcium fluxes.

| Calcium fluxes | Receptors | Molecules |
| --- | --- | --- |
| ECS to Cyto by VGCC | EAAT | Calcium |
| Cyto to ECS by NCX | AMPA | NE |
| Cyto to ECS by PMCA | mGluR5 (Gq) | Glutamate |
| ER to cyto by IP<sub>3</sub>R2 | mGluR3 (Gi) | Dopamine |
| ER to cyto by leakage | β-adrenergic (Gs) | Glutamine |
| Cyto to ER by SERCA | $\alpha_2$-adrenergic (Gi) | ROS |
| Mito to cyto by mPTP | $\alpha_1$-adrenergic (Gq) | ATP |
| Mito to cyto via membrane | D1-like (Gs) | PLC |
| Cyto to Mito via membrane | D2-like (Gi) | IP<sub>3</sub> |
|  | Heteromers of D1&D2 (Gq) | DAG |
|  | IP<sub>3</sub>R2 | cPKC |
|  |  | PKA |
|  |  | PDE |
|  |  | cAMP |
|  |  | Adenylyl cyclase |

Abbreviations: ECS: extracellular space; Cyto: cytoplasm; Mito: mitochondria.

**Complete model parameter list**

**Table 2.**
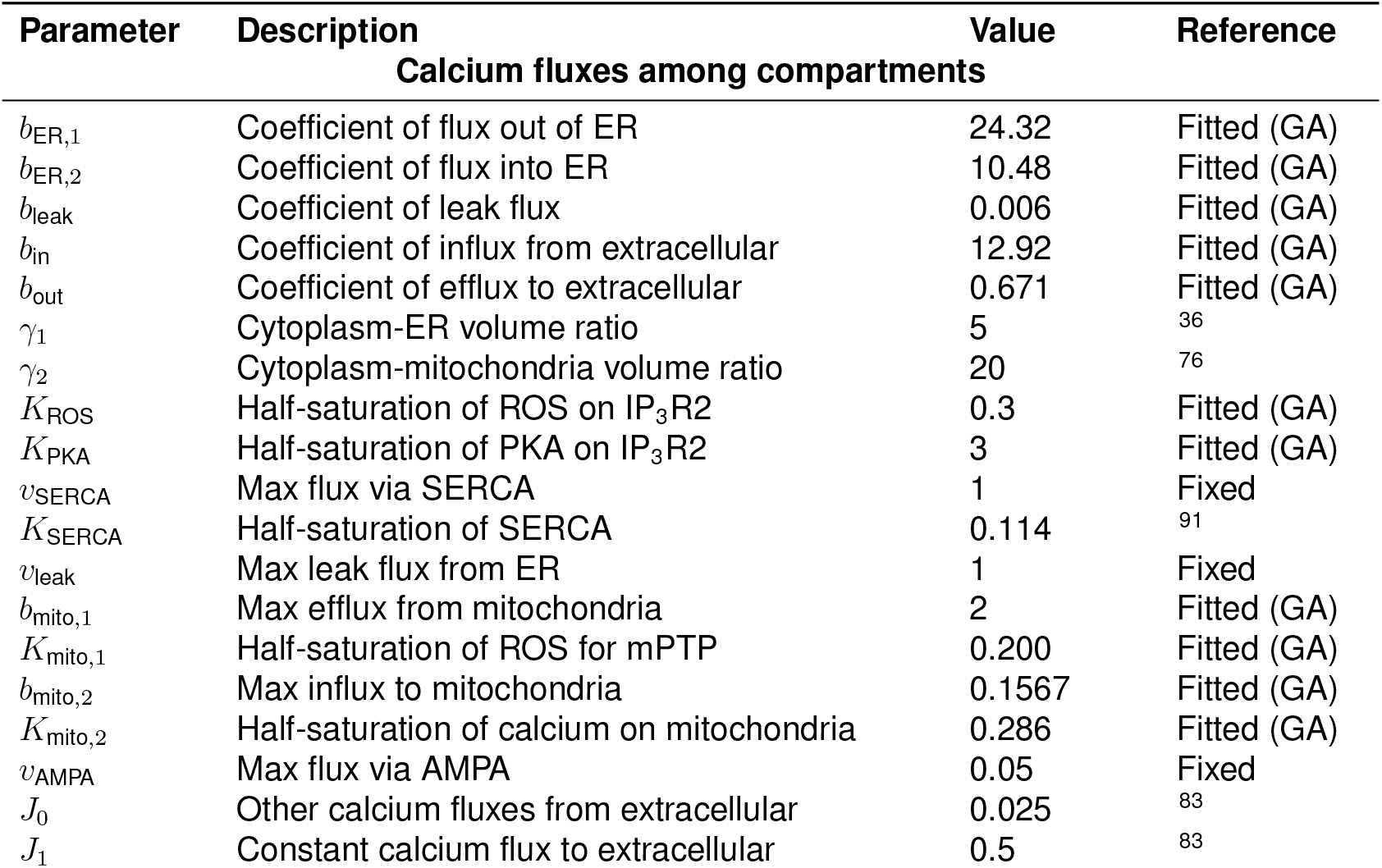
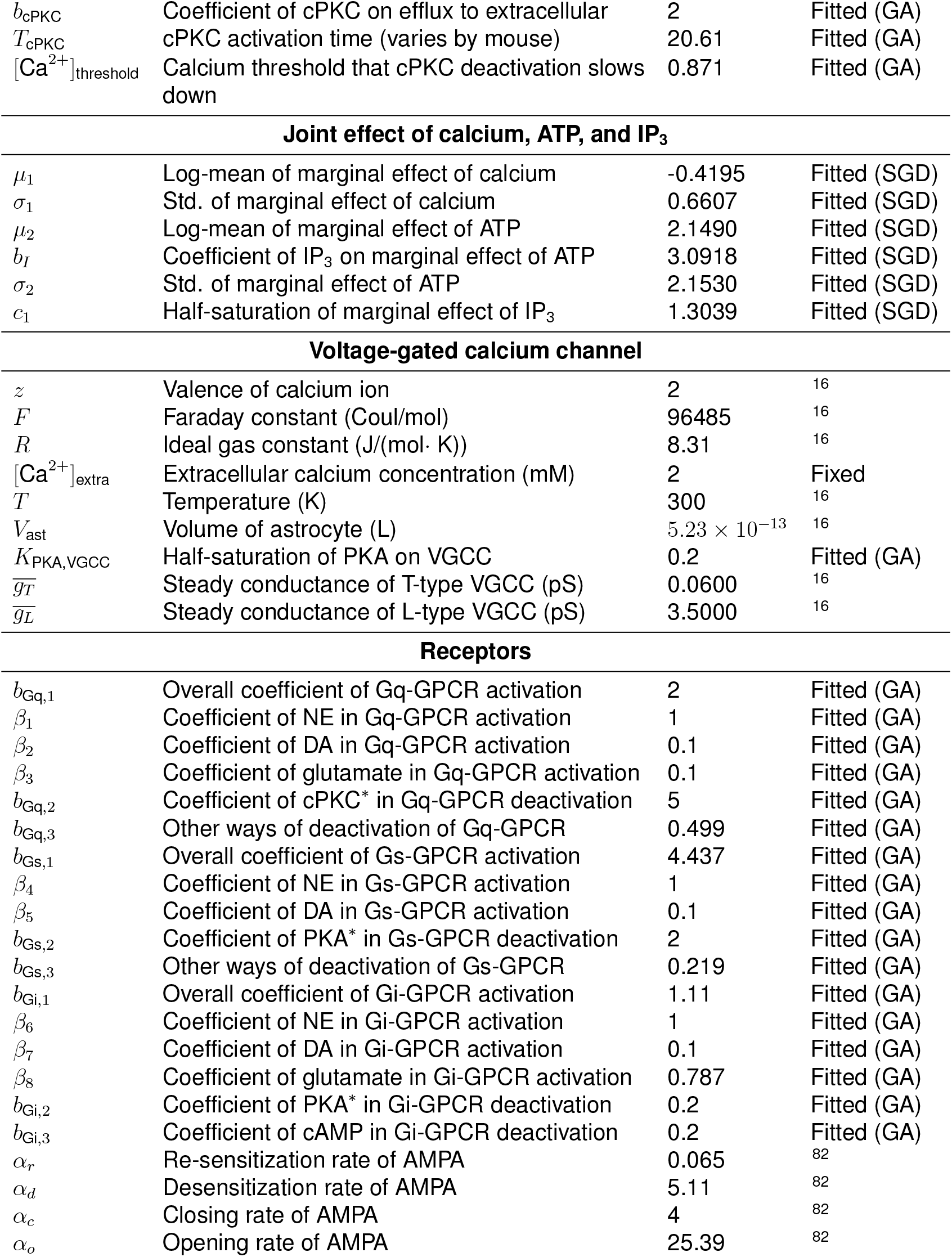
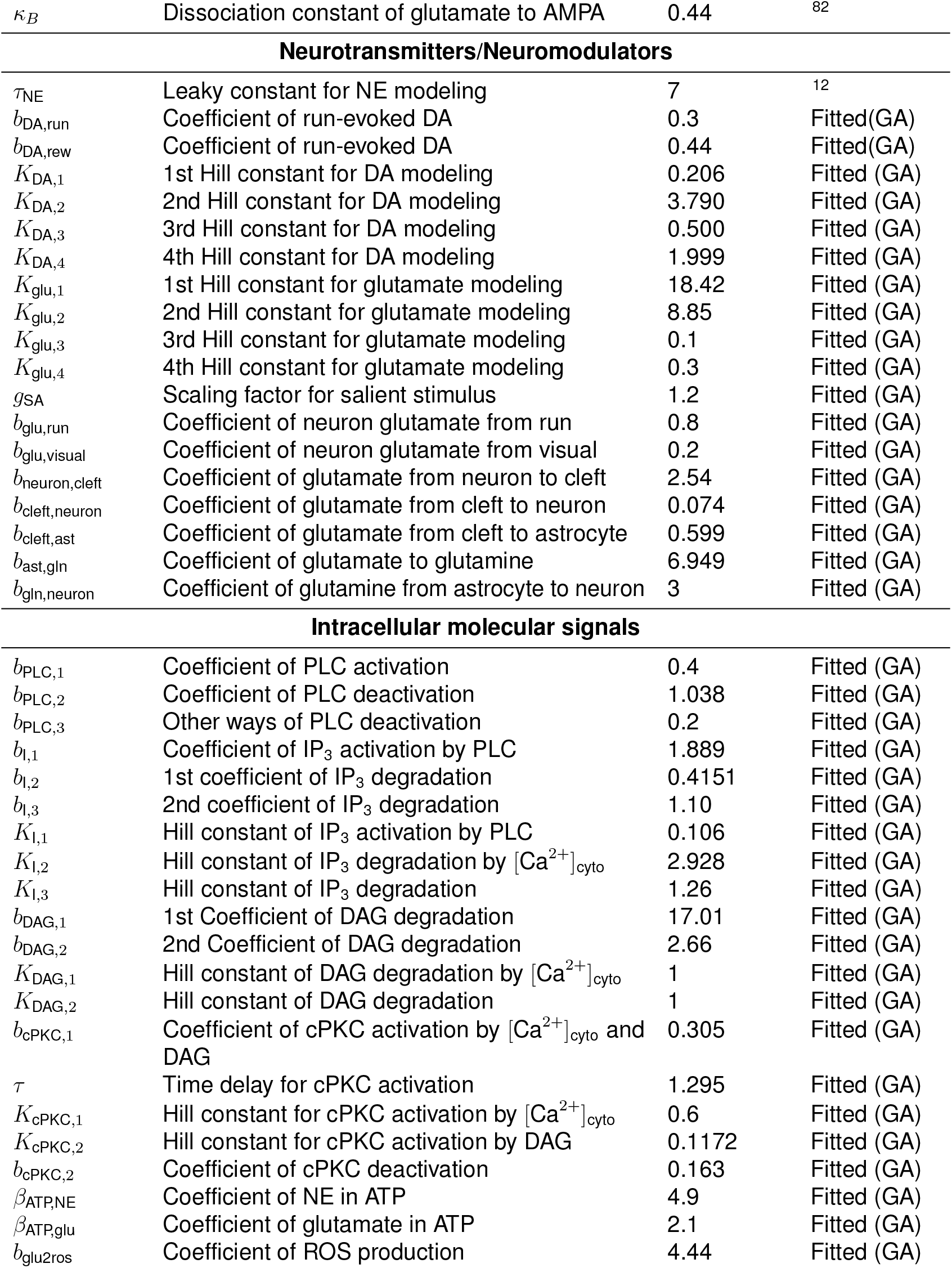
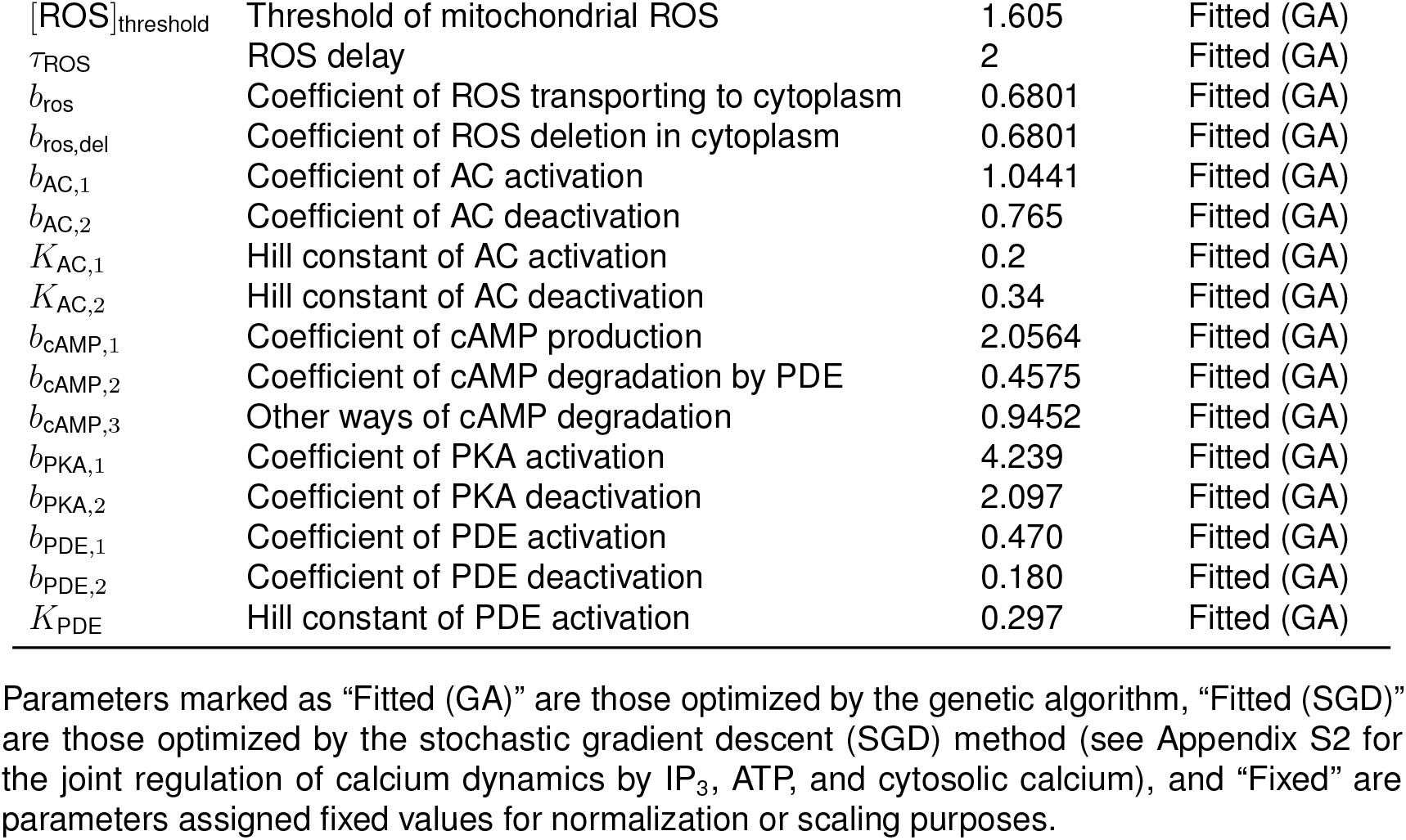
Parameter values and initial conditions.

## STAR METHODS

### Key resources table

**Table 3.** Key resources used in this study.

| REAGENT or RESOURCE | SOURCE | IDENTIFIER |
| --- | --- | --- |
| **Software and algorithms** |  |  |
| MATLAB R2023a | MathWorks | https://www.mathworks.com/products/matlab.html |
| Calcium signaling model | This paper | https://github.com/jameschengpeng/Calcium_model_explaining_refractory |
| AQuA2 (MATLAB version) | Mi *et al.*<sup>92</sup> | https://github.com/yu-lab-vt/AQuA2 |

### Method details for animal experiments

#### Experimental model and animal details

All procedures were conducted in accordance with the National Institutes of Health (NIH) guide for the Care and Use of Laboratory Animals and were approved by the Salk Institute Institutional Animal Care and Use Committee (IACUC). We employed GFAP-Cre 73.12 (RRID:IMSR JAX:012886) and Ai95D (RRID:IMSR JAX:024105) transgenic mouse lines. Behavioral and imaging experiments were performed using four heterozygous male mice. Two of them underwent two separate surgeries, a head-plate implant at 7–10 weeks of age and a cranial-window implant at ∼13 weeks of age, whereas the other two mice received a combined surgery at 8–12 weeks of age. The task-training started ∼7 days after each surgery. During training and imaging, mice were water-restricted to 25 ml·kg^−1^ per day to maintain 80–85% of their ad-libitum body weight. Optical recordings were performed between 20 and 26 weeks of age. The mice were group-housed with bedding and nesting materials under a 12 h light-dark cycle in a temperature (∼22-23^*°*^C) and humidity-controlled (∼45–65%) environment, with ad-libitum access to standard chow and water outside of experimental sessions.

#### Experimental timeline and study design

All animal subjects underwent several experimental procedures. Each mouse was first habituated for 1-2 days to minimize stress. All animals then received head-plate implantation and craniotomy surgery, followed by an approximately one-week recovery period. Two of the mice first received a head-plate implant, recovered, were habituated, and trained on the ball task prior to undergoing craniotomy surgery. After recovery from the craniotomy, these mice resumed task training. The remaining two mice underwent a combined procedure in which the head plate and craniotomy were performed in a single surgery before training began. All four mice successfully completed the behavioral task training and were subsequently imaged to measure astrocytic responses during task performance.

#### Surgical procedures and cranial window preparation

Surgical implantation of the head plate and cranial window was carried out as described previously^9,55,93^. Mice were anesthetized with isoflurane (4% for induction, 2% for maintenance) on a custom surgical bed (Thorlabs Inc., Newton, NJ), and body temperature was held at 36-37^*°*^C using a DC temperature controller. Ophthalmic ointment was applied to protect the eyes from drying The surgical site was cleaned with 70% ethanol and Betadine, and a midline skin incision of ∼10 mm was made. A section of the scalp was excised to reveal the frontal, parietal, and interparietal regions of the skull. A custom metal head plate was secured over the motor cortex using C&B Metabond Quick Adhesive Cement (Parkell Inc., Edgewood, NY), with cement extending to cover all other exposed areas of the skull.

A custom-made cranial window was implanted to permit long-term two-photon imaging^94^. For two mice, the window implantation was performed after recovery from head-plate surgery and initial behavioral training, whereas for the remaining mice, it was conducted during a single procedure immediately following head-plate implantation. The skull over the motor cortex was carefully thinned, and a circular craniotomy (2.5 mm diameter; centered at AP 1.5 mm, ML 1.5 mm) was made while keeping the dura mater intact. The craniotomy was sealed using a custom three-layer glass window assembly (each No. 1 thickness), with the two inner layers composed of 2.5 mm-diameter circular coverslips and the outermost layer made of a 3 mm-diameter circular cover glass resting on the thinned skull. The coverslips were bonded sequentially using UV-curable optical adhesive (NOA 71; Norland Products, Inc., Cat. No. 7106), ensuring the absence of trapped air bubbles that could impair imaging quality or lead to detachment of the cover glass during the postoperative period.

#### Behavioral setup and data acquisition

Animal training was conducted in a custom-built setup located within a sound-attenuating cubicle (ENV-017M, Med Associates Inc.). The setup featured a color LCD monitor for stimulus presentation (12.1” LCD Display Kit/500 cd/VGA, ICP Deutschland GmbH). To minimize optical recording noise, the monitor was covered with a color filter (R342 Rose Pink, Rosco Laboratories Inc.). The setup also included a spherical treadmill (Habitrail Mini Exercise Ball, Animal World Network), which allowed the animal to run either voluntarily or during task performance. Mice were positioned on the treadmill facing the LCD display, and head fixation was achieved by securing the head plate with custom-built holders. An optical encoder (E7P OEM, US Digital) attached to the treadmill measured both the speed and direction of ball rotation. Water rewards were delivered using a programmable syringe pump (NE-500 OEM Syringe Pump, New Era Pump Systems, Inc.). Behavior-related signals were collected via a data acquisition board (PCI-6221, National Instruments) connected to a breakout box (BNC-2110, National Instruments) and interfaced with MATLAB using the Data Acquisition Toolbox (Version R2010b SP2, The MathWorks Inc.). The behavioral task sequence was controlled by the MATLAB-based software MonkeyLogic (www.monkeylogic.net)^95,96^. Custom functions were integrated into MonkeyLogic to enable analysis and control of ball rotation parameters. Treadmill encoder signals and trial marker codes generated by MonkeyLogic were acquired at a 10 kHz sampling rate (±5 V input range) in synchronization with imaging data. Simultaneous acquisition using the microscope’s software (MScan; Sutter Instrument Company, 2016, 64-bit Version 2.2.0.0) allowed precise temporal alignment of run parameters, behavioral events, and image frames.

#### Behavioral task

Each trial began once the mouse remained stationary on the ball for 1 second. Initially, a blue square appeared on the monitor, during which the mouse needed to remain still for 20 seconds. If the mouse remained still throughout this period, a second stimulus (a filled blue square) appeared for 3 seconds in half of the trials, prompting the mouse to start running. This stimulus appeared at two intensity levels: either salient or near the perceptual detection limit, determined empirically late in training and kept constant during data collection. If the mouse began sustained running during the 3-second filled square stimulus presentation (ball speed exceeding 2 mm/s for at least 1 second), it received a water reward (classified as a Hit trial). If the mouse failed to run during this stimulus presentation (ball speed less than 0.5 mm/s), the trial was classified as a Miss trial In the other half of trials without a filled square stimulus, the mouse earned a water reward by remaining still for 3 seconds, which was classified as a Correct Rejection (CR) trial. If the mouse ran during this period, the trial was marked as a False Alarm (FA). Any movement by the mouse during the initial 20-second stand-still period (ball speed *>*2 mm/s) aborted the trial, categorizing it as a spontaneous (Spont) run. After a 5-second inter-trial interval (ITI), the mouse was able to initiate a new trial.

#### Animal habituation and behavioral training

Prior to behavioral training, mice were handled and tamed on two consecutive days to minimize stress. Training began with two days of habituation to the ball, during which mice were placed in the setup for approximately 15-30 minutes per day to become accustomed to both the ball and the head restraint. After habituation, mice remained on the ball for 60-90 minutes per day for 2-7 days to learn to balance and run. Only afterward were mice trained on the behavioral protocol. The training strategy and parameters were adjusted individually according to each mouse’s behavioral tendencies and rate of progress. Each daily training session lasted 60-90 minutes, during which mice performed approximately 300-700 trials per session. Behavioral performance and water reward were closely monitored to maintain a daily water intake of 25 ml/kg.

Training initially involved rewarding mice simply for running. Once a mouse reliably initiated 3-5 runs within a 5-minute window, visual stimuli were introduced with long presentation durations, and rewards were given only for runs initiated while the stimulus was visible. Over subsequent sessions, stimulus durations were gradually shortened and inter-stimulus intervals were lengthened. Once mice reached the performance criterion, stimuli were presented only after the mouse had stopped running and remained still for several seconds, with this stillness interval being progressively extended. In parallel, reward thresholds were adjusted to require specific acceleration, speed, and run duration parameters. Stimulus durations were further decreased, and reward amounts were fine-tuned to maintain motivation. When mice achieved full proficiency in the task, data acquisition began.

#### *In vivo* two-photon calcium imaging

The mice were imaged daily during task performance for 5-12 sessions. Imaging was performed using a resonant-scanning two-photon microscope (Sutter Instrument) equipped with a pulsed femtosecond Ti:Sapphire laser (Chameleon Ultra II, Coherent) for simultaneous optical and analog data acquisition. GCaMP6f fluorescence was excited at 910 nm and detected through an ET525/70M emission filter (Chroma Technology Corp.) and an H7422-40 GaAsP photomultiplier tube (Hamamatsu Photonics). The average excitation power varied with imaging depth and typically ranged from approximately 55 to 66 mW. The imaging depth was typically 100-135 *µ*m below the pia, corresponding to cortical layer 2/3^97^. Data were acquired using a Nikon 16×0.8 NA water-immersion objective. To minimize stray light from the LCD monitor, a custom blackout curtain was placed around the microscope’s detector. Images (512 × 512 pixels) were collected at 1.0×Zoom, yielding an effective field of view of approximately 510 *µ*m × 640 *µ*m after cropping, at a frame rate of ∼30.9 frames/s using MScan software (Sutter Instrument Company, 2016, 64-bit Version 2.2.0.0).

Each imaging session consisted of five to twelve recordings of approximately 10 minutes each, separated by short breaks of 3-5 minutes. Recordings within the same session were acquired at the same cortical site to maximize the number of trial repetitions for subsequent analysis. Across sessions, conducted on consecutive days, recording sites were shifted laterally or axially to increase the total sampled tissue volume.

#### Image and behavior data preprocessing

To maintain real-time control over task-related events during data collection, running speed was analyzed using brief, unprocessed measurements. A detailed post hoc review of the velocity patterns and recorded trial types was carried out, and only those trials that matched the protocol-required criteria were included in the final dataset. All analyses were conducted using MATLAB scripts specifically developed for this purpose. The system translated voltage change frequencies from the encoder into running speeds, which were then processed with an exponential filtering technique. This filtering method retained essential velocity variations while minimizing noise, allowing for the reliable detection of running periods and event durations. The filtered velocity was calculated iteratively using a weighted combination of current and previous values.

To address timing shifts introduced by the velocity filter, a 0.2-second tolerance window was applied when aligning data to key task events, such as trial or reward onset. A velocity threshold of 0.5 mm/s was set to define the beginning and end of a run. This low threshold was chosen because even minimal motion could trigger calcium activity, and it was important to confirm the absence of movement during stationary periods and during the stimulus phase in CR or Miss trials. Any run beginning within 0.5 seconds of the end of a prior run was considered part of the same running event. For a run to be classified as part of a Hit or FA trial, it needed to last more than 1 second and exceed a speed of 10 mm/s. If the ball remained stationary (below 0.5 mm/s) during the stimulus period, the trial was marked as a CR trial. Unlike Hit and FA trials, which consistently included a run during stimulus presentation, there was no run during the stimulus phase for Miss and CR trials. Nonetheless, Miss and CR were often followed by a run shortly after the end of the stimulus or following the delivery of water reward, respectively.

### Quantification and statistical analysis

#### Data structure and preprocessing pipeline

We recorded several calcium videos from each mouse. Each video covers a field of view (FOV) of approximately 510 *µ*m × v640 *µ*m, with ∼22,000 frames, recorded at a sampling rate of ∼30.9 frames per second (fps). The spatial resolution was ∼1 *µ*m per pixel, and the FOV contained approximately 80 astrocytes based on histology. Each recording included multiple trials. For each calcium video, we simultaneously acquired mouse run-velocity data and timestamps of visual stimuli or reward delivery.

#### Time-lapse calcium image data analysis

As reported in the literature^98^, intracellular calcium events in astrocytes typically last hundreds of milliseconds, with most occurring on a timescale of seconds or tens of seconds. Therefore, the ∼30.9 fps temporal resolution of our calcium videos is more than sufficient for analyzing astrocyte calcium activity. To optimize data handling, we downsampled the calcium videos in the temporal domain by averaging every 10 frames. This improved signal-to-noise ratio while preserving the temporal resolution required to capture calcium transients.

To further improve the signal-to-noise ratio, we downsampled in the *X* and *Y* dimensions by a factor of 2 using bicubic interpolation in MATLAB. In our data, this spatial downsampling presents an acceptable compromise between computational efficiency and minimizing information loss. Astrocytes, including the soma and processes, span an approximately 50 × 50*µ*m territory; hence, this downsampling does not risk losing signal from astrocyte processes. We validated this method against alternative approaches, such as bilinear interpolation, and found minimal impact on signal fidelity.

After spatial and temporal downsampling, we employed AQuA2^92^, an event-based software designed for quantifying calcium activity in time-lapse imaging data. Unlike traditional region-of-interest (ROI)-based tools, AQuA2 automatically detects independent calcium events, where each calcium event is a group of voxels that are spatially connected and exhibit a single temporal peak, without the need for predefined ROIs, a limitation in traditional analysis tools. This feature allows AQuA2 to capture calcium dynamics with varying spatial distributions over time, overcoming the limitations of fixed ROIs. Figure S1 shows examples of calcium events detected by AQuA2.

By analyzing the calcium events detected by AQuA2 across all calcium recordings, we found that calcium events triggered by rewarded trials occupied a larger number of pixels compared to non-rewarded trials. An unpaired two-sample t-test performed on the number of pixels occupied by calcium events in both trial categories yielded a 95% confidence interval for the difference between population means of [1636.5, 2847.2] with a p-value of 5.203 × 10^−13^ (Figure S1G). Additionally, since the distributions of pixel counts for both types of trials exhibited heavy tails, we used log-transformed values on the vertical axis of Figure S1. The empirical cumulative distribution functions (CDFs) for both types of trials reveal that the CDF curve for rewarded trials consistently lies beneath that of non-rewarded trials, indicating a heavier-tailed pixel count distribution for rewarded trials (Figure S1H). Taken together, these results indicate that rewarded trials elicit broader, larger-scale calcium activities across the spatial domain than non-rewarded trials at the population level (Figure S1G, H).

By examining the rising time maps of calcium events across all recordings, defined as the time at which each pixel in an event reaches 50% of its amplitude, we generated spatial summaries of early-responding pixels (Figure S1I, J). Figure S1I shows a normalized heatmap from an example recording, highlighting pixels that consistently responded early during calcium events. Pixels with values ≥ 0.5 were classified as frequent early-response pixels in the normalized heatmaps Figure S1J shows the clustered early-activation density (CEAD) scores across all recordings on the population level. The CEAD score is defined as the proportion of frequent early-response pixels within the FOV. A smaller CEAD score indicates more condensed early-response regions Our results indicate that early-response regions in each recording are relatively condensed, with no more than 25% of pixels (most below 10%) responding early following a stimulus (Figure S1I, J). Consequently, we compiled the reference calcium signal for each recording by averaging the calcium signals from the largest early-response region.

